# Centromeric chromatin clearings demarcate the site of kinetochore formation

**DOI:** 10.1101/2024.04.26.591177

**Authors:** Kathryn Kixmoeller, Yi-Wei Chang, Ben E. Black

## Abstract

The centromere is the chromosomal locus that recruits the kinetochore, directing faithful propagation of the genome during cell division. The kinetochore has been interrogated by electron microscopy since the middle of the last century, but with methodologies that compromised fine structure. Using cryo-ET on human mitotic chromosomes, we reveal a distinctive architecture at the centromere: clustered 20-25 nm nucleosome-associated complexes within chromatin clearings that delineate them from surrounding chromatin. Centromere components CENP-C and CENP-N are each required for the integrity of the complexes, while CENP-C is also required to maintain the chromatin clearing. We further visualize the scaffold of the fibrous corona, a structure amplified at unattached kinetochores, revealing crescent-shaped parallel arrays of fibrils that extend >1 μm. Thus, we reveal how the organization of centromeric chromatin creates a clearing at the site of kinetochore formation as well as the nature of kinetochore amplification mediated by corona fibrils.

## Main Text

Human centromeres are defined by the presence of nucleosomes containing the histone H3 variant CENP-A (Earnshaw and Rothfield, 1985; Kixmoeller et al., 2020; Musacchio and Desai, 2017). An individual kinetochore contains multiple kinetochore protein complexes which assemble on CENP-A nucleosomes and form a link between centromeric chromatin and the microtubules of the mitotic spindle. CENP-A nucleosomes recruit the inner kinetochore which is composed of the 16-subunit constitutive centromere associated network (CCAN) (Foltz et al., 2006; Kixmoeller et al., 2020; Musacchio and Desai, 2017; Okada et al., 2006), present at the centromere throughout the cell cycle. During mitosis, the inner kinetochore recruits the outer kinetochore which is primarily responsible for binding to the microtubule-based spindle (Kixmoeller et al., 2020; Musacchio and Desai, 2017). Multiple copies of the inner and outer kinetochore protein complexes work as an assembly to form the functional kinetochore. During prometaphase and states of chromosome misalignment, the kinetochore further recruits the fibrous corona, a complex assembly of proteins, including fibrils of the ROD–Zwilch–ZW10 (RZZ) complex with Spindly, which serves to activate the spindle assembly checkpoint and capture microtubules (Kops and Gassmann, 2020; McAinsh and Kops, 2023; Raisch et al., 2022).

Early electron microscopy of the kinetochore in cells revealed a trilaminar structure with a structured inner kinetochore plate at the surface of the chromatin, an unstructured electron-translucent region, and a structured outer kinetochore plate (Brinkley and Stubblefield, 1966; Rieder, 1982). Beyond the outer plate, a crescent-shaped fibrous region, the fibrous corona, was present only on kinetochores not attached to microtubules (Brinkley and Stubblefield, 1966; Jokelainen, 1967; Rieder, 1982). Later experiments using improved sample preparation techniques alongside immunofluorescence approaches revealed that the apparent rigid trilaminar structure of the kinetochore was influenced by traditional dehydrated sample preparation and suggested instead a less-rigid, mesh-like structure for the kinetochore (Blower et al., 2002; Dong et al., 2007; Magidson et al., 2015; McEwen et al., 1998; McIntosh et al., 2013). Recent structural studies of *in vitro* reconstituted inner kinetochore components have shown the CCAN to be a globular complex closely associated with a CENP-A nucleosome (Pesenti et al., 2022; Tian et al., 2022; Yatskevich et al., 2022). On the other hand, the outer kinetochore consists of narrow, elongated protein complexes (Alushin et al., 2010; Kixmoeller et al., 2020; Musacchio and Desai, 2017; Nishino et al., 2013; Petrovic et al., 2016; Valverde et al., 2016) which may contribute to the less-structured region located between the inner kinetochore and the fibrous corona or area of microtubule binding.

At the level of kinetochore-forming centromeric chromatin, CENP-A-containing nucleosomes are thought to be found in multiple patches along the linear DNA sequence interspersed with canonical nucleosomes (i.e., those harboring conventional histone H3) (Blower et al., 2002; Ribeiro et al., 2010; Zinkowski et al., 1991). These proposals are supported and extended by genomic approaches that build on the recent complete human centromere assemblies (Altemose et al., 2022a; b; Dubocanin et al., 2023; Gershman et al., 2022; Logsdon et al., 2021).

Importantly, the human kinetochore has never been directly visualized at molecular resolution in its chromosomal context, as *in situ* approaches have until now been limited to fluorescence-based studies (Blower et al., 2002; Magidson et al., 2015) or the aforementioned electron microscopy studies of fixed samples (Brinkley and Stubblefield, 1966; Dong et al., 2007; Magidson et al., 2015; McEwen et al., 1998; Rieder, 1982). In contrast, cryo-electron tomography (cryo-ET) permits 3D visualization of a vitreous sample of interest. This approach captures proteins in their native conformations and natural arrangements, unperturbed by chemical fixation and heavy metal staining during sample preparation which obscures molecular details especially at the level of chromatin. In this way, we reasoned that cryo-ET would be capable of visualizing the human kinetochore in its native context on isolated chromosomes to reveal individual kinetochore complexes and their organization both along the DNA strand and in 3D space.

### The kinetochore revealed by cryo-ET

The small size of kinetochore-forming centromeric chromatin relative to intact mitotic chromosomes necessitated the use of correlative light and electron microscopy (CLEM) to locate the kinetochore within isolated chromosomes for cryo-ET data collection. A typical human interphase centromere is estimated to contain 100-400 copies of the CENP-A protein, and thus 50-200 CENP-A nucleosomes (Bodor et al., 2014). CENP-A nucleosomes are split between sister chromatids during DNA replication in S-phase, so in the subsequent mitosis each sister kinetochore harbors only 25-100 CENP-A nucleosomes, out of ∼650,000 total nucleosomes in an average-sized human chromosome. In the DLD-1 cells used in our cryo-ET experiments, centromeres have CENP-A nucleosomes which number at the low end of this range (Bodor et al., 2014).

Isolated mitotic chromosomes have previously been imaged by cryo-ET to investigate bulk chromatin on chromosome arms and demonstrate the irregular packing of mitotic chromatin (Beel et al., 2021). For kinetochore studies, biochemical isolation of mitotic chromosomes has historically provided many foundational insights for the field (i.e., to understand central aspects such as microtubule nucleation (Telzer et al., 1975), poleward movement on microtubules (Koshland et al., 1988), and generation of a diffusible spindle checkpoint signal (Kulukian et al., 2009)), owing to the faithful retention of inner and outer kinetochore complexes. We isolated mitotic chromosomes from DLD-1 cells in which CENP-A is biallelically tagged at its endogenous locus with rsEGFP2 to allow for CLEM (Fig. 1A, S1A,B).

**Figure 1:**
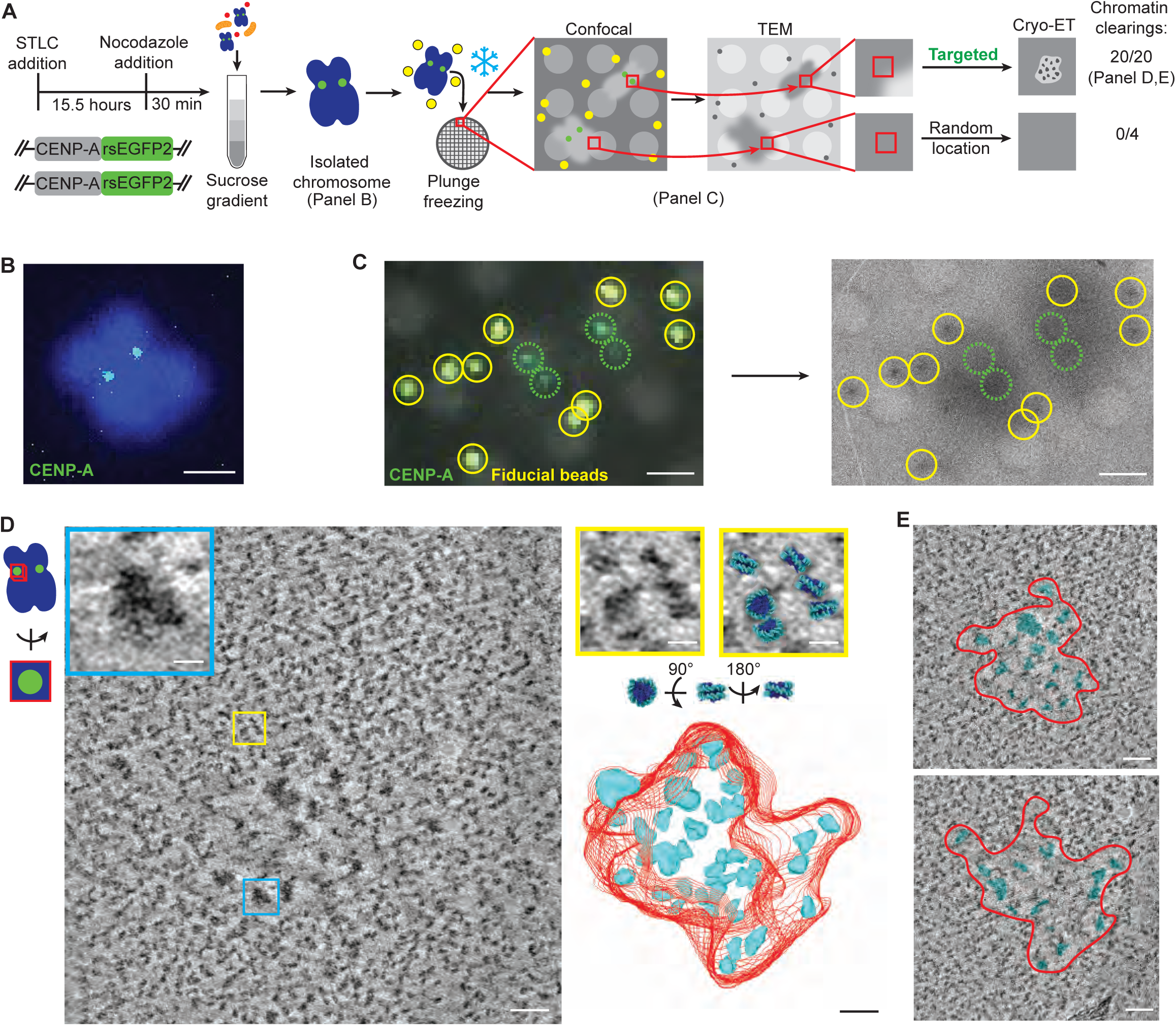
Targeting and visualizing the chromatin architecture of the kinetochore. **A.** Schematic of the experimental approach including mitotic chromosome isolation, EM grid preparation, CLEM, and the incidence of chromatin clearings in tomograms targeted to CENP-A fluorescence or to random locations on chromosome arms. An example tomogram acquired at a random location on the chromosome arm is shown in Fig. S5B. **B.** Example of an isolated and partially decondensed mitotic chromosome with intact morphology and CENP-A fluorescence at sister kinetochores, imaged on a coverslip. Scale bar, 2 μm. **C.** Example of CLEM: two chromosomes on EM grids are shown by both cryogenic confocal microscopy (left) and transmission electron microscopy (right). Yellow circles mark fluorescent beads. Green dashed circles highlight the location of kinetochores in both modalities. Scale bars, 2 μm. **D.** Example tomogram slice showing kinetochore complexes within a chromatin clearing located at the chromosome surface (left, also see Fig. S2) and 3D model of the full locus (right). This tomogram was obtained from a different chromosome than those shown in (C) and the kinetochore is oriented such that the chromatin surface is roughly parallel to the page, as depicted in the cartoon to the left. In the model, the boundary of the chromatin clearing is traced in red and kinetochore complexes are traced in cyan. Cyan inset shows an example kinetochore complex. Yellow inset shows a region of chromatin outside the kinetochore with an overlay of nucleosome structures (PDB: 1KX3). Scale bars, 50 nm (insets 10 nm). **E.** Overlay of the model with other slices of the same tomogram. Scale bars, 50 nm.

Isolated mitotic chromosomes retained expected morphology and the characteristic “double dot” pattern of CENP-A fluorescence representing the sister kinetochores (Fig. 1B). Both inner kinetochore and outer kinetochore components are retained on isolated chromosomes, with no detectable loss during the steps prior to grid preparation (Fig. S1C,D). Grids containing vitreous chromosomes and 200 nm fluorescent fiducial beads (Fig. 1C, yellow foci) were imaged by cryogenic confocal microscopy and kinetochores were identified by paired “double dot” green fluorescent foci (Fig. 1C). Grids were then transferred to a cryogenic transmission electron microscope for cryo-ET data collection. The 200 nm fluorescent beads, visible by both confocal and transmission electron microscopy, were used to carry out correlation between the two imaging modalities (Schorb and Briggs, 2014; Schellenberger et al., 2014) (Fig. 1A,C). In this way, our approach allowed us to collect tilt series at kinetochores with high targeting accuracy (Fig. 1A).

In tomograms reconstructed from these tilt series, we consistently identified a distinctive chromatin architecture located at the surface of the chromosome (Fig. 1A,D,E, S2,S3, Videos S1,S2). Within the dense, irregularly packed nucleosomes that comprise the majority of mitotic chromatin (Beel et al., 2021; Chen et al., 2023), we found chromatin clearing(s) containing larger proteinaceous densities which we hypothesized to be inner kinetochore complexes (Fig. 1A,D,E, S2,S3). The chromatin clearings are distinct from the surrounding chromatin and have a well-defined boundary (traced in red, Fig. 1D,E) which segregates kinetochore complexes from surrounding chromatin. Within chromatin clearings, these larger densities are separated by stretches of open area which significantly exceed the separation between nucleosomes in surrounding chromatin (Fig. 2A). We find that the majority of kinetochores (65%) contain a single chromatin clearing (e.g., Fig. 1D, Video S1), but a single kinetochore may also be comprised of 2 (25%) or 3 (10%) separate chromatin clearings (e.g., Fig. S3A and Video S2: two clearings coalescing at the chromosome surface, Fig. S4A: two separated clearings). Kinetochore chromatin clearings contain fewer particles per unit area than surrounding chromatin (Fig. 2B,C, S5A). The particles within chromatin clearings are also larger on average (Fig. 2B,D, S5A) and further separated from neighboring particles (Fig. 2B,E, S5A) than in the surrounding chromatin. Chromatin clearings were identified in all tomograms targeted to CENP-A fluorescent foci and were not identified in tomograms collected at random chromosome locations (Fig. 1A, S5B).

**Figure 2:**
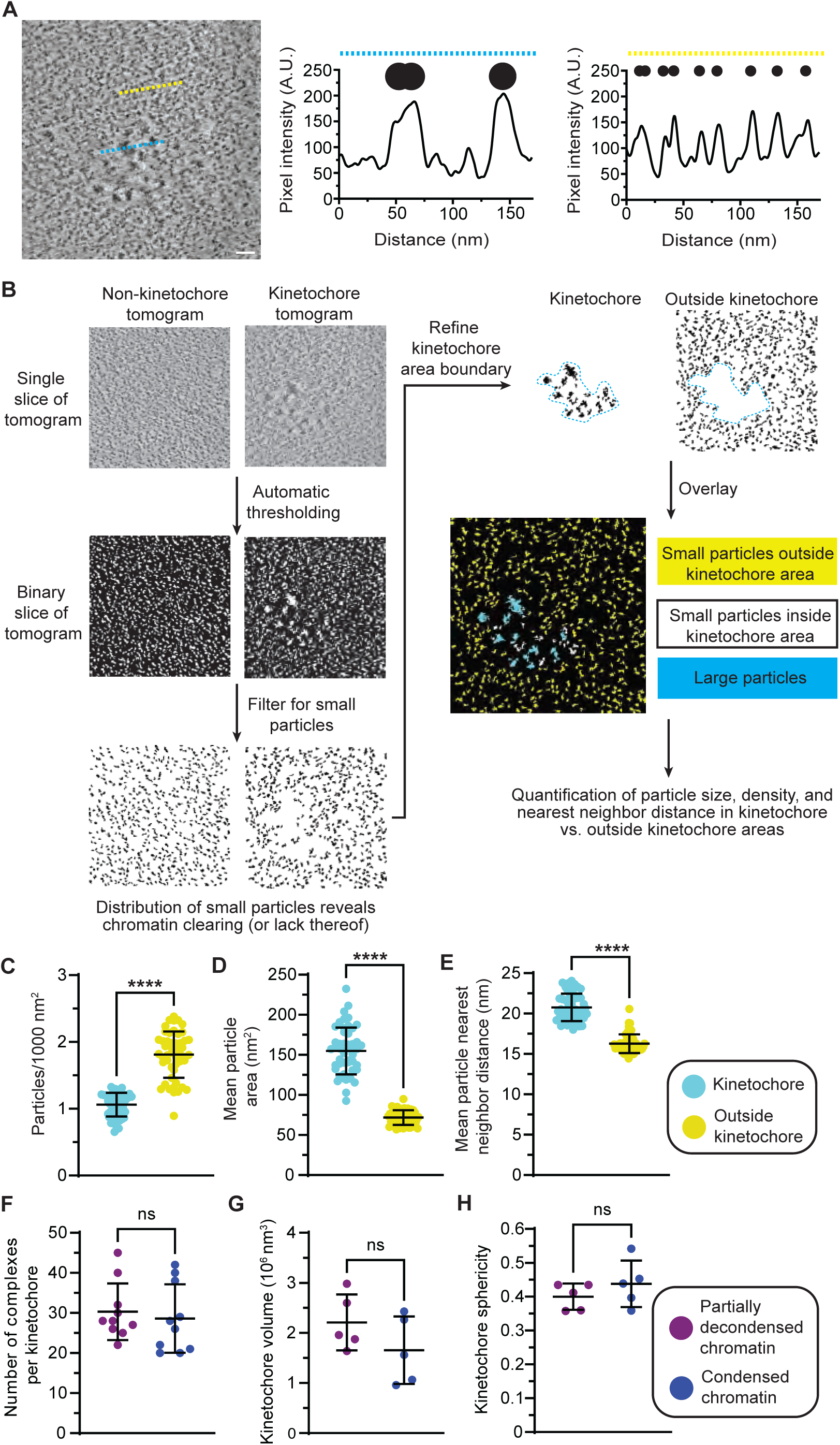
The inner kinetochore consists of 20-25 nm particles in a clearing devoid of dense chromatin. **A.** Line plots highlighting the extent of clearing around complexes within kinetochore area. Left: tomogram slice of an example kinetochore (from Fig. 1D) with the lines used to generate line plots through the kinetochore area (cyan) and surrounding chromatin (yellow). Right: Line plots and cartoon representation of the density profile. Higher pixel intensity values represent darker pixels. Within the kinetochore region (cyan), larger densities (kinetochore particles) are punctuated by long stretches of empty space (clearing). In contrast, surrounding chromatin consists of smaller, more tightly packed densities (nucleosomes) with only small spacing between particles. Line plots were generated after Gaussian filtering (σ = 2.0) of the tomogram slice. **B.** Schematic of the workflow used to quantify kinetochore architecture for (C-E). The overlay image highlights agreement between the automatic filtering for small particles (e.g., nucleosomes) and the manually-refined kinetochore area. The kinetochore area contains mostly large particles (cyan) and relatively few small particles/nucleosomes (white). **C-E.** Quantitative comparison of kinetochore versus nearby chromatin. Data are presented as mean ± SD. **C.** Particles per 1000 nm^2^: 1.06 ± 0.18 vs. 1.81 ± 0.35 (N = 50, p < 0.0001, t = 17.73, df = 49). **D.** Mean particle area (nm^2^): 154.9 ± 29.26 vs. 71.78 ± 9.10 (N = 50, p < 0.0001, t = 23.53, df = 49). **E.** Mean particle nearest neighbor distance (nm): 20.77 ± 1.70 vs. 16.28 ± 1.16 (N = 50, p < 0.0001, t = 16.45, df = 49). **F-H.** Quantitative comparison of kinetochores from partially decondensed chromatin versus condensed chromatin. **F.** Number of complexes per kinetochore: 30.30 ± 7.07 vs. 28.60 ± 8.53 (N = 10, p = 0.63, t = 0.49, df = 18). **G.** Kinetochore volume (10^6^ nm^3^): 2.21 ± 0.56 vs. 1.66 ± 0.67 (N = 5, p = 0.19, t = 1.42, df = 8). **H.** Kinetochore sphericity: 0.40 ± 0.04 vs. 0.44 ± 0.07 (N = 5, p = 0.31, t = 1.08, df = 8). Note that sphericity provides a quantitative assessment of 3D shape, but there is no assumption or expectation that a kinetochore be spherical.

In terms of the copy number of inner kinetochore complexes, we find that an individual kinetochore contains on average 30 larger complexes (Fig. 2F), consistent with estimates for the number of CENP-A nucleosomes at DLD-1 kinetochores (Bodor et al., 2014). The number of complexes does vary among kinetochores (Fig. 2F), consistent with findings that CENP-A occupancy varies among centromeres even within an individual (Altemose et al., 2022a; Gershman et al., 2022). This architecture is also seen in chromosome preparations with more condensed chromatin (Fig. S4), and the number of kinetochore complexes observed, the volume of the kinetochore area, and the sphericity of the kinetochore are similar between partially decondensed and condensed chromatin preparations (Fig. 2F-H). We conclude that the large-scale architecture of the kinetochore is preserved in our preparations but it is impossible to eliminate the possibility of some distortion from blotting onto EM grids. For these reasons, we note that our main focus is on the smaller-scale architecture and details of kinetochore complexes. In sum, the chromatin clearings on the chromosome surface revealed by our cryo-ET studies spatially separate the relatively large, dense kinetochore complexes from the rest of the chromosome.

### Inner kinetochore complexes

Close inspection of the larger densities found within chromatin clearings (Fig. 3A, Video S3) reveal a variety of densities, the most prevalent of which are globular particles measuring 20-25 nm in their longest dimension and roughly triangular in shape (Fig. 3B), consistent in size and shape with inner kinetochore CCAN complexes. These densities lack the relatively high degree of structural homogeneity seen in bulk nucleosomes (Beel et al., 2021)(Fig. 3B), indicating that the protein-protein and/or protein-DNA interfaces within them permit local flexibility rather than existing as highly rigid bodies on the chromosome surface. Nucleosomes are visible embedded within these complexes (Fig. 3C, S6A,B), identified by density corresponding to two gyres of DNA with spacing and curvatures consistent with wrapping an octameric histone core. Since the CCAN is associated with CENP-A-containing nucleosomes(Foltz et al., 2006), we conclude that the nucleosomes closely packing with CCAN densities contain CENP-A. We note that some proposed models (Dalal et al., 2007; Pesenti et al., 2022) include a centromeric histone particle harboring only a single wrap of DNA on a core with only a single copy of CENP-A and core histones H2A, H2B, and H4 (i.e., a hemisome). Our finding of nucleosomes with two gyres of DNA visible (Fig. 3C, S6A,B) support other models in which the CCAN is bound to an octameric CENP-A-containing nucleosome (Foltz et al., 2006; Hasson et al., 2013; Pesenti et al., 2022; Yatskevich et al., 2022). At the level of individual inner kinetochore complexes, our experiments reveal that centromeric DNA is linked to the kinetochore by close association of the CCAN complex with an octameric nucleosome.

**Figure 3:**
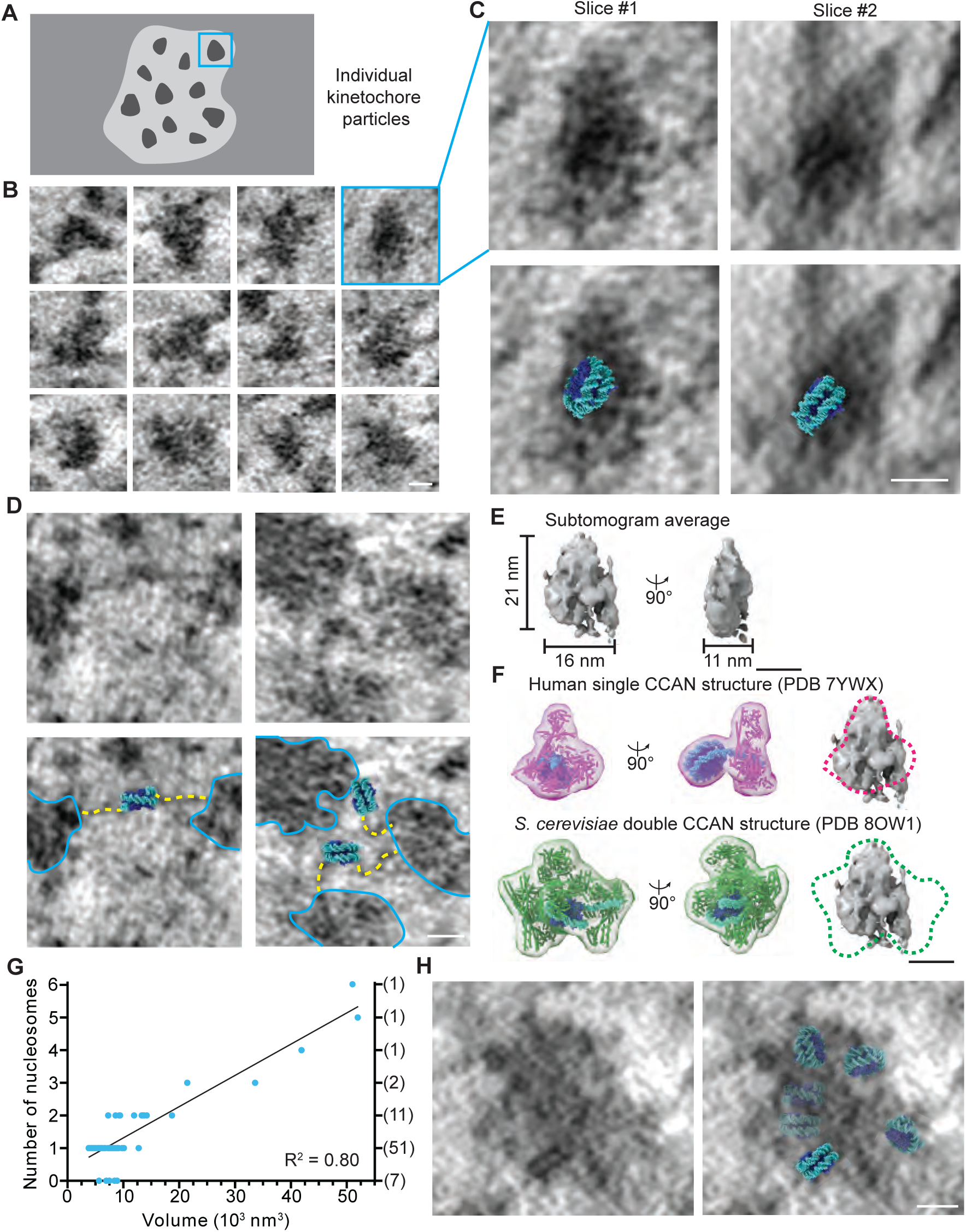
Kinetochore complexes harbor closely associated nucleosomes and are typically separated by a linker nucleosome. **A.** Cartoon depicting closer examination of a kinetochore complex within a chromatin clearing. **B.** Multiple examples of individual kinetochore complexes seen in tomogram slices. Scale bars, 10 nm. **C.** An example kinetochore complex viewed from two angles. Densities corresponding to the double gyres of an octameric nucleosome are visible embedded within the kinetochore complex. Lower panels show overlay with nucleosome structure. **D.** Two examples of connections observed between adjacent kinetochore complexes, annotated in lower panels. Kinetochore complexes, or multimers thereof (right panel), are outlined in cyan, and yellow dashed lines trace DNA strands. **E.** Subtomogram average of kinetochore complexes demonstrates the size and shape of individual inner kinetochore complexes. **F.** Comparison of published structures of the reconstituted inner kinetochore from human and yeast (*S. cerevisiae*) to kinetochore complexes in our tomograms. Left: two published structures of the inner kinetochore. In magenta is the human single CCAN with CENP-A nucleosome (Yatskevich et al., 2022)(PDB: 7YWX). In green is the yeast (*S. cerevisiae*) inner kinetochore with two copies of the CCAN flanking a CENP-A nucleosome (Dendooven et al., 2023)(PDB: 8OW1). Each structure is shown within a 50 Å resolution envelope. Right: envelopes of the reconstituted inner kinetochore structures overlaid with the subtomogram average of kinetochore complexes from our tomograms. **G.** Comparison of kinetochore particle volume versus number of nucleosomes identified within the density shows a linear relationship. The number of particles for each nucleosome number is shown in parentheses (right). The few particles in which no nucleosome is detected likely contain a nucleosome in an orientation that is unfavorable for visualization. **H.** Example tomogram slice showing a larger multimer with one embedded nucleosome clearly visible. The rough positions of the other 5 nucleosomes identified within this density (in other tomogram slices) are shown as partially transparent in the overlay (right).

Connections between adjacent kinetochore complexes, consisting of linker DNA and a single intervening nucleosome, were readily observable within our tomograms (Fig. 3D). The intervening nucleosome lacks a kinetochore complex and in some cases is located very close to one of the two complexes (Fig. 3D right panel, S6B). This finding suggests that kinetochore complexes are clustered along stretches of DNA and often assemble on alternating nucleosomes. The kinetochore complexes within a given chromatin clearing may all be positioned on a contiguous stretch of DNA or may represent multiple shorter contiguous regions of kinetochore occupancy, separated on the linear DNA molecule by stretches of conventional nucleosomes, but in a manner that coalesces in three dimensions.

We next investigated the size and shape of the inner kinetochore complexes to compare with existing models built on reconstitutions from purified components (Dendooven et al., 2023; Pesenti et al., 2022; Tian et al., 2022; Yatskevich et al., 2022). As we predicted, the biological complexity captured in our tomograms of the kinetochore precluded straightforward structural characterization. Kinetochore complexes contain flexible components (Pesenti et al., 2022; Tian et al., 2022; Yatskevich et al., 2022) and are able to interact with a mix of binding partners (Kixmoeller et al., 2020; Malvezzi et al., 2013; Rago et al., 2015), leading to a structurally heterogeneous population. Furthermore, the complexes are few in number and embedded in a crowded chromatin environment (to an extent not found in many other cellular structures, if at all), factors which seriously hinder subtomogram averaging efforts (Fig. S7A,B), whereas canonical nucleosomes were more easily averaged (Fig. S7C,D). For these reasons, our goal was not to produce a high-resolution structure of the kinetochore complex. However, we did note that the complexes were fairly uniform in size and shape (Fig. 3B), permitting us to measure their average dimensions (∼16 × 21 × 11 nm^3^) (Fig. 3E, S7A,B). Since CENP-A nucleosomes contain two copies of CENP-A, each of which contains potential binding sites for CCAN complex recruitment, models exist for either one or two copies of the CCAN per CENP-A nucleosome (Allu et al., 2019; Weir et al., 2016). We reasoned that the size of the particles in our tomograms could distinguish between these models. Recent structures of the *in vitro* reconstituted human inner kinetochore contain a single copy of the CCAN per CENP-A nucleosome (Fig. 3F, upper panel; S7B)(Yatskevich et al., 2022), but the equivalent structure of the *Saccharomyces cerevisiae* inner kinetochore contains two copies of the CCAN complex flanking a single CENP-A nucleosome (Fig. 3F, lower panel; S7B)(Dendooven et al., 2023). To distinguish between models with one versus two copies of the CCAN per particle, we compared the shapes of these two inner kinetochore structures to our subtomogram average (Fig. 3E,F, S7B) as well as to individual particles from our tomograms (Fig. 3B). We found that the structure containing two copies of the CCAN is too large to be consistent with the kinetochore particles visualized in our tomograms, whereas a single copy of the CCAN is a closer match in size and shape (Fig. 3F, S7B). Therefore, our cryo-ET studies indicate that the repeating unit of the human inner kinetochore is a single copy of the CCAN complex associated with a CENP-A nucleosome.

In addition to the individual inner kinetochore particles, larger globular densities are also observed (Figs. 1D and 3G). These larger densities were also found to contain nucleosomes and the number of nucleosomes scales linearly with the volume of the density (Fig. 3G), suggesting that these larger particles represent higher-order packing of inner kinetochore complexes into multimers. These multimers vary from close association of two complexes (Fig. S6C) to much larger multimers containing up to 6 nucleosomes (Fig. 3H, S6D). We note that the vast majority of inner kinetochore complexes are monomeric (Fig. 3G), but along with the relatively rare multimer assemblies of nucleosome-harboring inner kinetochore complexes, they represent the major structured element of the inner kinetochore that can be visualized via cryo-ET.

### Architectural role for CENP-C

The chromatin clearing around the CCAN complexes that helps to separate the inner kinetochore from bulk chromatin was an unexpected finding in our tomograms. We predicted that the CCAN component CENP-C would contribute to this architecture since it resembles a molecular “scaffold” (Carroll et al., 2010; Klare et al., 2015; Tanaka et al., 2009). More specifically, CENP-C is an elongated protein which binds many kinetochore components (Carroll et al., 2010; Klare et al., 2015; Tanaka et al., 2009), dimerizes through its C-terminal domain (Cohen et al., 2008; Hara et al., 2023; Trazzi et al., 2009), binds a second nucleosome (Klare et al., 2015; Hara et al., 2023; Ali-Ahmad et al., 2019; Kato et al., 2013; Walstein et al., 2021), and maintains the integrity of centromeric chromatin upon physical stretching (Ribeiro et al., 2010). These properties could allow CENP-C to organize not only individual CCAN complexes but also the kinetochore as a whole. To test the effects of acute CENP-C removal, we employed a DLD-1 cell line derivative in which CENP-C is biallelically tagged with eYFP and an auxin inducible degron (AID) tag (Fig. 4A)(Fachinetti et al., 2015; Guo et al., 2017; Hoffmann et al., 2016). In this cell line, addition of the synthetic auxin indole-3-acetic acid (IAA) leads to rapid degradation of CENP-C to undetectable levels within 30 minutes (Guo et al., 2017). We isolated chromosomes from this cell line and collected cryo-ET data as before, with and without the addition of IAA for an hour prior to chromosome isolation (Fig. 4B). To detect kinetochores for CLEM on purified chromosomes where CENP-C-eYFP is removed, we used fluorescent antibody labeling of CENP-A during the chromosome purification protocol (Fig. 4B). In tomograms collected after degradation of CENP-C (Fig. 4B, lower panel; 4D, S8A), but not in those collected without CENP-C degradation (Fig. 4B, upper panel; 4C, S8B), we noted a pattern of disruption to kinetochore architecture with a spectrum of severity (Fig. 4D,E, S8A). Although some kinetochores retained the normal chromatin architecture of distinct kinetochore complexes within a chromatin clearing, consistent with the resiliency of the kinetochore complex against disruptions (Hoffmann et al., 2016), the majority of tomograms showed a partial or total loss of this architecture (Fig. 4D,E). In many cases, the chromatin clearing is lost, with individual kinetochore complexes still identifiable but with greatly reduced separation between adjacent kinetochore complexes and between the kinetochore complexes and the surrounding chromatin (Fig. 4D, left panel). In other cases, we observed a more severe phenotype in which kinetochore architecture is greatly perturbed. The kinetochore can still be identified as a region which differs from surrounding chromatin, but the individual kinetochore complexes are indistinct, and the boundaries between the kinetochore and surrounding chromatin are poorly defined (Fig. 4D, right panel; Video S4). These experiments provide evidence that CENP-C contributes to organizing the architecture of the kinetochore and specifically to the distinctive chromatin clearing around kinetochore complexes revealed by our cryo-ET data.

**Figure 4:**
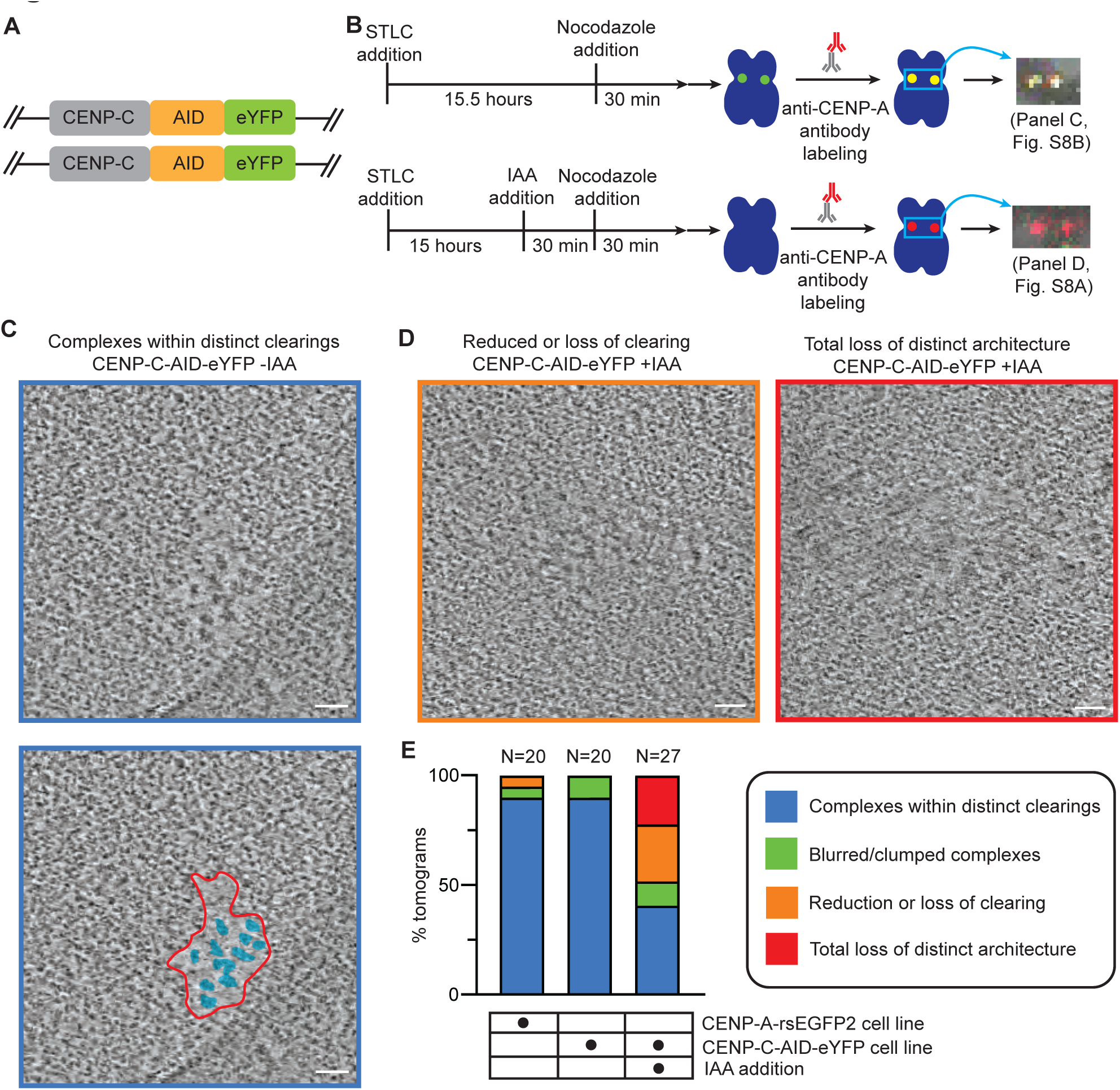
Removal of CENP-C perturbs kinetochore architecture. **A.** Diagram of the CENP-C-AID-eYFP cell line used for these experiments. **B.** Schematic of the experimental approach. Cryo-confocal images are shown of example sister kinetochores from each condition; images are overlay of green, far-red, and T-PMT channels. **C.** Representative image showing normal kinetochore morphology (complexes within distinct chromatin clearing) from CENP-C-AID-eYFP -IAA condition. Annotated in lower panel, showing the outline of chromatin clearing in red and kinetochore complexes in cyan. Note an edge of the underlying carbon film grid is visible in the bottom center of the image. Scale bars, 50 nm. **D.** Representative images from CENP-C-AID-eYFP +IAA condition showing kinetochores with reduction or loss of chromatin clearing (left, orange), and total loss of distinct architecture (right, red). **E.** Incidence of each kinetochore morphology in three experimental conditions: CENP-A-rsEGFP2 (N=20, initial dataset), CENP-C-AID-eYFP -IAA (N=20), and CENP-C-AID-eYFP +IAA (N=27).

### CENP-N supports CCAN integrity, not chromatin clearings

If the nature of CENP-C is required for chromatin clearing at the kinetochore (Carroll et al., 2010; Klare et al., 2015; Tanaka et al., 2009), then manipulations that retain CENP-C but disrupt the CCAN are predicted to leave the clearings intact. CENP-N is a central component of the CCAN (Pesenti et al., 2022; Tian et al., 2022; Yatskevich et al., 2022; Allu et al., 2019; Carroll et al., 2010; McKinley et al., 2015), and, importantly, its degradation in mitosis destabilizes other CCAN components without impacting CENP-C localization (McKinley et al., 2015)(Fig. S9A). We isolated chromosomes from a cell line expressing CENP-N-eGFP-AID under these conditions (Fig. 5A,B) and used fluorescent antibody labeling of CENP-C (Bassett et al., 2010) during the chromosome purification protocol to detect kinetochores for CLEM on purified chromosomes where CENP-N-eGFP is removed (Fig. 5B). In tomograms collected after degradation of CENP-N, we observed a striking decrease in the size of the kinetochore complexes (Fig. 5C-I, S9B,C, Video S5) which include closely associated nucleosomes (Fig. 5F-H). We quantified the volumes of these smaller densities, as in Fig. 3G, and found them to be ∼30% the size of the native inner kinetochore complexes (Fig. 5I). This magnitude of this decrease in volume, coupled with the continued presence of nucleosomes in these densities, lead us to conclude that the smaller densities are most consistent with a CENP-A nucleosome bound to CENP-C. This conclusion is in agreement with previous work demonstrating that other CCAN components are lost after mitotic degradation of CENP-N (McKinley et al., 2015). Since CENP-C directly interacts with outer kinetochore complexes (Malvezzi et al., 2013; Rago et al., 2015; Przewloka et al., 2011; Gascoigne et al., 2011; Screpanti et al., 2011), the more proximal elements of the outer kinetochore likely also contribute to nearby residual density after CENP-N depletion. We note that other nearby density remaining after CENP-N degradation is typically fibrillar in nature and not of the appearance of a nucleosome-associated complex (Fig. 5D, S9B, Video S5). Importantly, despite the evident perturbation of the inner kinetochore after CENP-N degradation, the smaller kinetochore complexes remain surrounded by a distinct chromatin clearing (Fig. 5D-J, S9B, Video S5), suggesting that the loss of CENP-N perturbs the CCAN while leaving the chromatin clearing architecture of the kinetochore intact. Thus, while it is possible that CENP-C is assisted by other factors in organizing the centromere, the chromatin clearings that distinguish this part of the chromosome do not require an intact CCAN.

**Figure 5:**
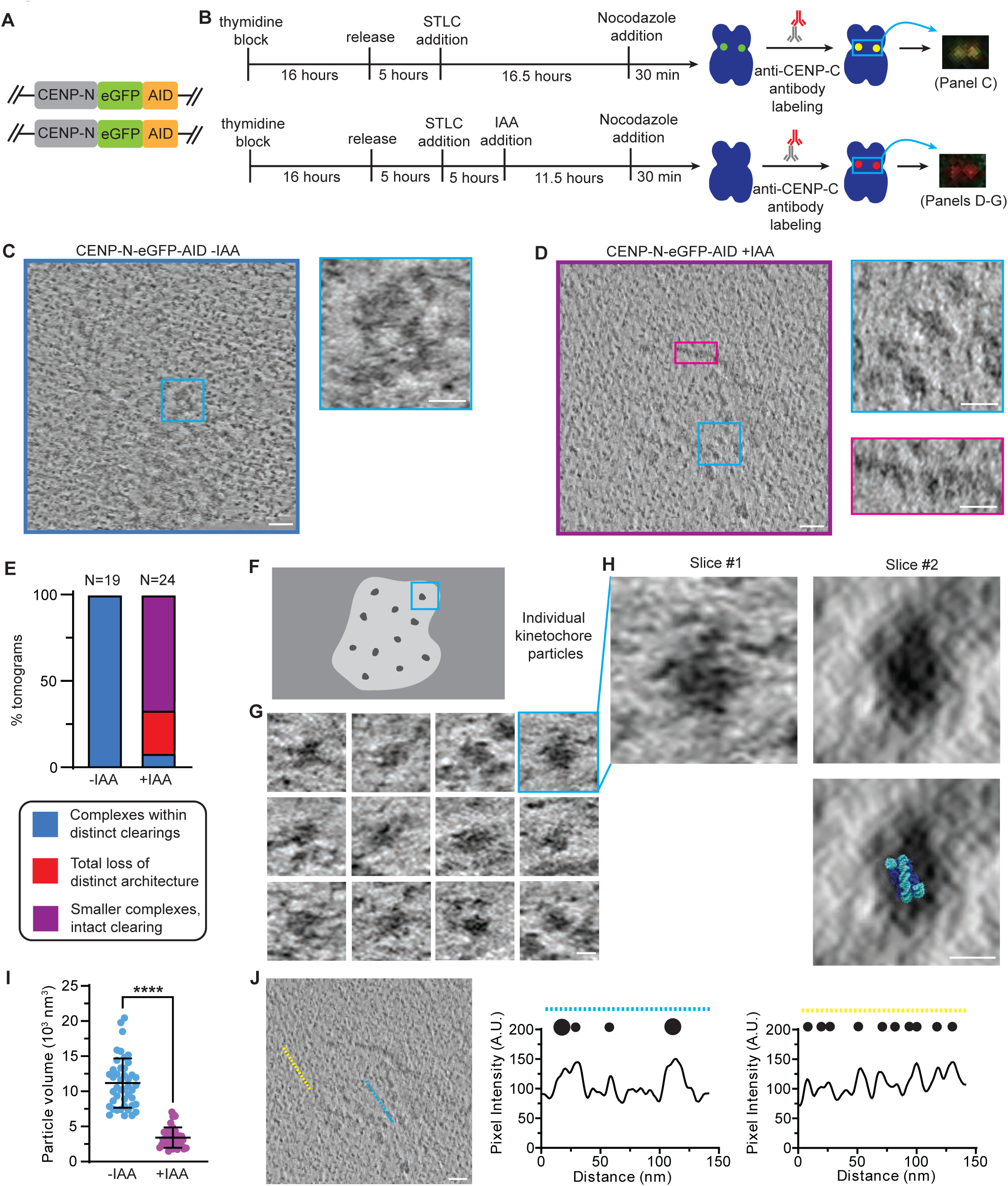
Removal of CENP-N perturbs inner kinetochore but leaves chromatin clearing intact. **A.** Diagram of CENP-N-eGFP-AID cell line used for this experiment. **B.** Schematic of the experimental approach. Cryo-confocal images are shown of example sister kinetochores from each condition; images are overlay of green and red channels. **C.** Representative image showing normal kinetochore morphology (complexes within distinct chromatin clearing) from CENP-N-eGFP-AID -IAA condition. Scale bar, 50 nm (Inset 20 nm). **D.** Representative image from CENP-N-eGFP-AID +IAA condition showing a kinetochore with kinetochore complexes that are notably reduced in size (cyan inset) but with chromatin clearing intact. Fibrous corona components are retained in this condition (magenta inset) but appear disorganized. Scale bar, 50 nm (Insets 20 nm). **E.** Incidence of each kinetochore morphology in the two experimental conditions: CENP-N-eGFP-AID -IAA (N = 19), CENP-N-eGFP-AID +IAA (N = 24). **F.** Cartoon depicting closer examination of a reduced-size kinetochore complex within a chromatin clearing. **G.** Multiple examples of reduced-size individual kinetochore complexes seen in tomogram slices. Scale bar, 10 nm. **H.** An example reduced-size kinetochore complex viewed from two angles. Densities corresponding to the double gyres of an octameric nucleosome are visible embedded within the kinetochore complex. Lower panel shows overlay with nucleosome structure. Scale bar, 10 nm. **I.** Quantification of kinetochore particle volume in the two experimental conditions, demonstrating that kinetochore particles are greatly reduced in volume after CENP-N degradation. Particle volume (10^3^ nm^3^): 11.17 ± 3.52 vs. 3.41 ± 1.45 (N = 82, p < 0.0001, t = 12.71, df = 80) **J.** Line plots highlighting the retained chromatin clearing around the reduced-size complexes within the kinetochore area. Left: tomogram slice of an example kinetochore (from Fig. 5D) with the lines used to generate line plots through the kinetochore area (cyan) and surrounding chromatin (yellow). Right: Line plots and cartoon representation of the density profile. Higher pixel intensity values represent darker pixels. Line plots were generated after Gaussian filtering (σ = 2.0) of the tomogram slice. Scale bar, 50 nm.

### Visualizing the fibrous corona

Beyond the inner kinetochore, the other part of the kinetochore that is predicted to have dense protein structure is the fibrous corona (Brinkley and Stubblefield, 1966; Jokelainen, 1967; Rieder, 1982). Whereas the inner kinetochore is thought to be relatively static in composition during mitosis, the fibrous corona is highly dynamic. The fibrous corona grows on the micron scale at kinetochores which are perpetually unattached from the spindle (Jokelainen, 1967; Kops and Gassmann, 2020). This growth is essential for generating a large microtubule binding surface to “rescue” the unattached chromosomes as well as to amplify the spindle assembly checkpoint so that a single unattached kinetochore can biochemically arrest the entire cell prior to the metaphase-to-anaphase transition (McAinsh and Kops, 2023). Thus, we anticipated that our cryo-ET experiments could clarify the architecture of this key structure. Indeed, in tomograms from our original dataset (Figs. 1-3; and noted, above, as being a retained feature after CENP-N depletion [Fig. 5D]), we observed the presence of short fibrous structures near the chromosome surface and in the vicinity of inner kinetochore complexes (Fig. 6A,B) which we hypothesized to be components of the fibrous corona. While relatively short, the fibrils are consistent in diameter with a key corona component, fibrils formed by the ROD-Zwilch-ZW10 (RZZ) complex in association with Spindly (Raisch et al., 2022).

**Figure 6:**
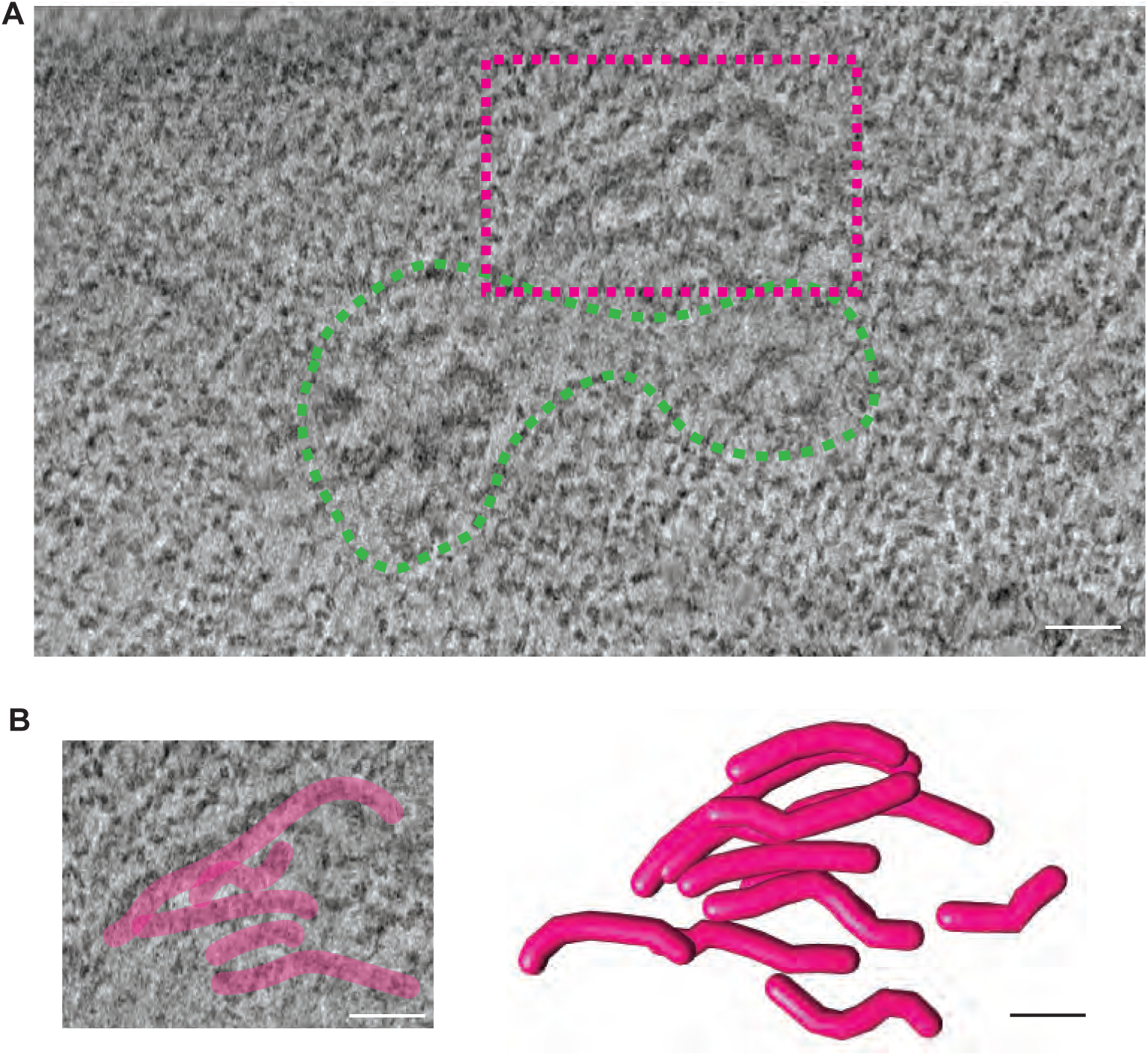
Corona fibrils observed near inner kinetochore complexes. **A.** Example tomogram slice showing kinetochore complexes in a chromatin clearing (green dashed outline) and a small fibrous corona (pink dashed outline). Scale bar, 50 nm. **B.** Inset of the fibrous corona from (A) with tracing of the corona fibrils visible in this tomogram slice (upper panel) and a model of the fibrils present in the full tomogram volume (lower panel). Scale bar. 50 nm.

If the fibrils we observed indeed represent the corona, then they should greatly expand when chromosomes are perpetually unattached from the spindle. We next tested this prediction. In our original preparations (Figs. 1-3), chromosomes experience only brief detachment from the mitotic spindle and so lack a large corona (Fig. 7A, upper panel; S10A)(Cooke et al., 1997). We modified our protocol to include prolonged treatment with the microtubule depolymerizer nocodazole such that chromosomes are persistently unattached from spindle microtubules to increase the size of the corona (Fig. 7A, lower panel with CENP-E as the marker of corona expansion (Cooke et al., 1997); Fig. S10B). Consistent with our prediction, we observed a marked expansion of the fibrous corona in tomograms collected at the kinetochores of chromosomes isolated after prolonged spindle detachment. We observe interwoven fibrils forming an extended mesh-like structure on a micron scale (Fig. 7B,C, S10C, Video S6). While interwoven, the fibrils are largely aligned on their long axis (Fig. 7B,C, S10C), indicating that they do not typically intersect in a perpendicular manner (e.g., as in nuclear lamina fibers). Individual fibrils are about 15 nm in width (Fig. 7D) and are consistent in dimension and appearance with fibrils of RZZ-Spindly formed *in vitro* and imaged by negative-stain electron microscopy (Fig. 7E)(Raisch et al., 2022). This supports the conclusion that RZZ-Spindly fibers are the dominant structural component of the fibrous corona, and we noted that interspersed among the RZZ-Spindly fibrils are other densities likely representing the other proteinaceous components of the corona. We find that the RZZ-Spindly fibrils arc around the inner kinetochore forming a crescent shape. The fibrils either wrap around a single kinetochore on its outward aspect or take on a more extended form bridging two sister kinetochores (Fig. 7F, S10C), matching corona profiles seen in mitotic cells (Fig. S10B). In its expanded state, individual RZZ-Spindly fibrils are not only longer in length, they also occasionally form larger “sheets”, appearing to be organized via lateral interactions among individual fibrils (Fig. 7G). Indeed, the lateral interactions appear central to generating a robust scaffold for the expanded fibrous corona. Taken together, our experiments reveal the arrangement of the fibrous corona of the kinetochore in its native context on intact chromosomes.

**Figure 7:**
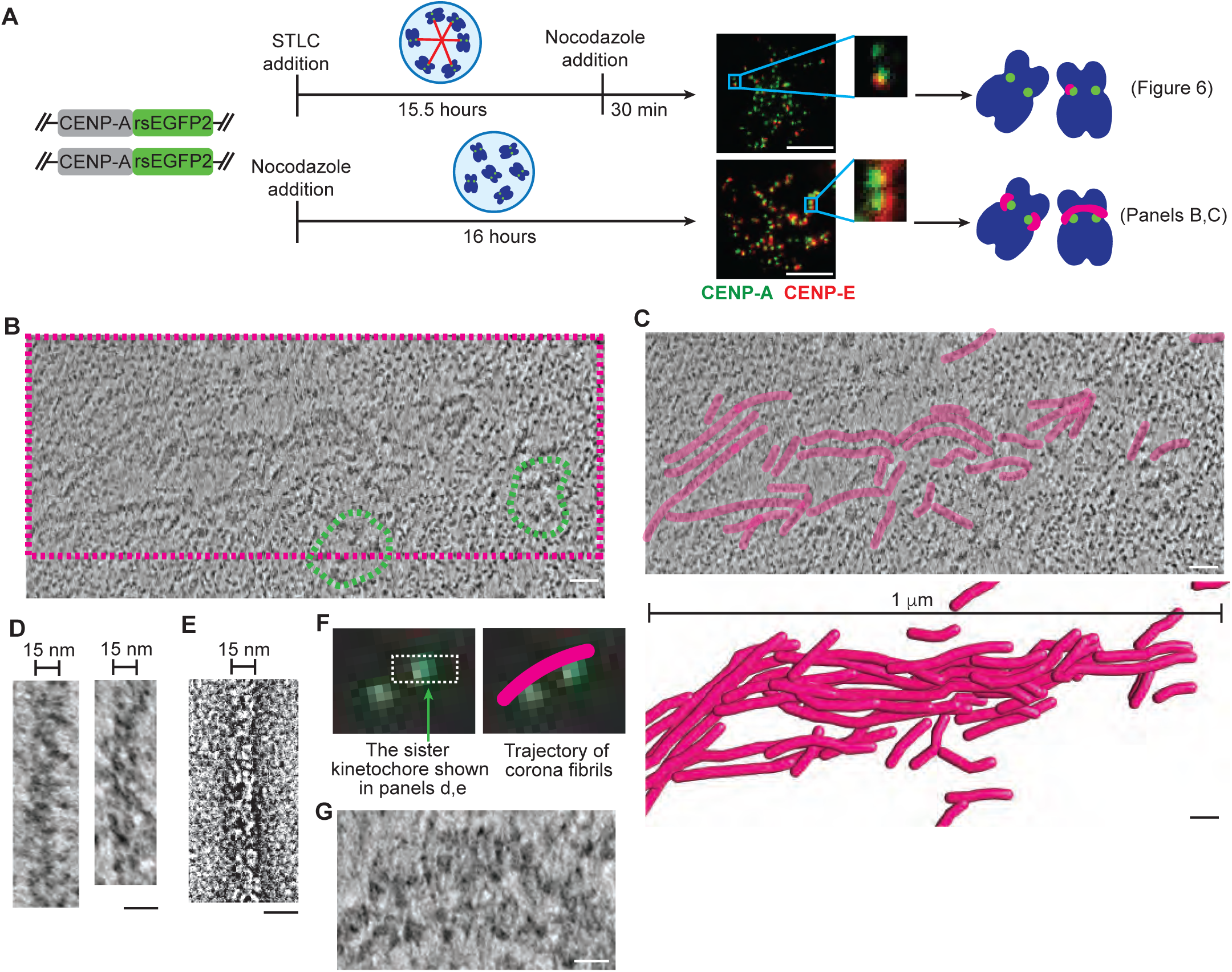
The scaffold of the fibrous corona consists of parallel ∼15 nm fibers that can extend >1 μm. **A.** Diagram of CENP-A-rsEGFP2 cell line used for this experiment and simplified schematic of the experimental approach. Upper panel shows original protocol, lower panel shows prolonged nocodazole treatment. Immunofluorescence images show occupancy of CENP-E, a corona component, at kinetochores under both conditions. **B.** Example tomogram slice from a chromosome isolated after prolonged nocodazole treatment showing kinetochore complexes in two chromatin clearings (green dashed outlines) and a large fibrous corona extending ∼1 μm in length (pink dashed outline). Scale bar, 50 nm. **C.** Inset of the fibrous corona from (B) with tracing of the corona fibrils visible in this tomogram slice (upper panel) and a model of the fibrils present in the full tomogram volume (lower panel). Scale bar, 50 nm. **D.** Two examples of corona fibrils in tomogram slices from chromosomes isolated after prolonged nocodazole treatment. Scale bar, 20 nm. **E.** For comparison, a negative-stain electron microscopy image of *in vitro* formed fibril of RZZ-GFP with farnesylated Spindly (BioImage Archive: S-BIAD364)(Raisch et al., 2022) which is consistent in dimension and appearance with the fibrils visualized in our tomograms (D). Scale bar, 20 nm. **F.** Cryo-confocal image of the chromosome imaged in (B) with the field of view from (B) outlined in white (left). Right: depiction of the trajectory of the corona fibrils shown in (B,C) beyond the tomogram field of view, bridging over to the other sister kinetochore. **G.** Inset from (B) showing the formation of sheet-like structures from apparent lateral interactions among corona fibrils. Scale bar, 20 nm.

## Discussion

Our cryo-ET studies reveal that the human kinetochore consists of clustered kinetochore complexes within distinctive chromatin clearings located at the chromosome surface. Two main structured features with clear densities emerge: the chromatin-linked inner kinetochore and the fibrous outer kinetochore corona. The kinetochore complexes visualized in our tomograms are consistent with a single copy of the human CCAN associated with an octameric CENP-A nucleosome. We provide evidence that the CCAN directly dictates its spatial delineation from the rest of the chromosome, as removing a key CCAN component, CENP-C, rapidly compromises the clearing that separates the inner kinetochore from nearby densely packed nucleosomes. In contrast, removing CENP-N causes reduction in the size of the kinetochore complexes while retaining the chromatin clearing. This structural foundation is required for downstream regulatory components (e.g., Bub1 kinase) and microtubule binding components (e.g., the Ndc80 complex) of the outer kinetochore that likely do not themselves directly contribute to the overall architecture. Cryo-ET also permitted visualization of the fibrous structures of the corona and its expansion on the surface of mitotic chromosomes upon prolonged spindle detachment.

These findings establish an updated architectural model of the human kinetochore (Fig. 8). Individual CCAN complexes are closely associated with a CENP-A-containing nucleosome. On the linear DNA, these CCAN complexes are often separated by linker DNA and a single nucleosome that is not directly associated with the CCAN. In three dimensions, however, the CCAN complexes come close together but exhibit striking spatial separation from the densely packed conventional chromatin that comprises the rest of the mitotic chromosome. The necessity of CENP-C in preserving this spatial separation aligns with earlier proposals suggesting its importance in kinetochore maintenance (Hara et al., 2023; Klare et al., 2015; Walstein et al., 2021), yet it is certainly not the only component of the kinetochore that may support the overall architecture. In our model, CENP-C links CCAN complexes to nucleosomes both inside and outside the chromatin clearing (Fig. 8) forming a web of inter-connections which hold together the kinetochore as a distinct domain within mitotic chromatin.

**Figure 8:**
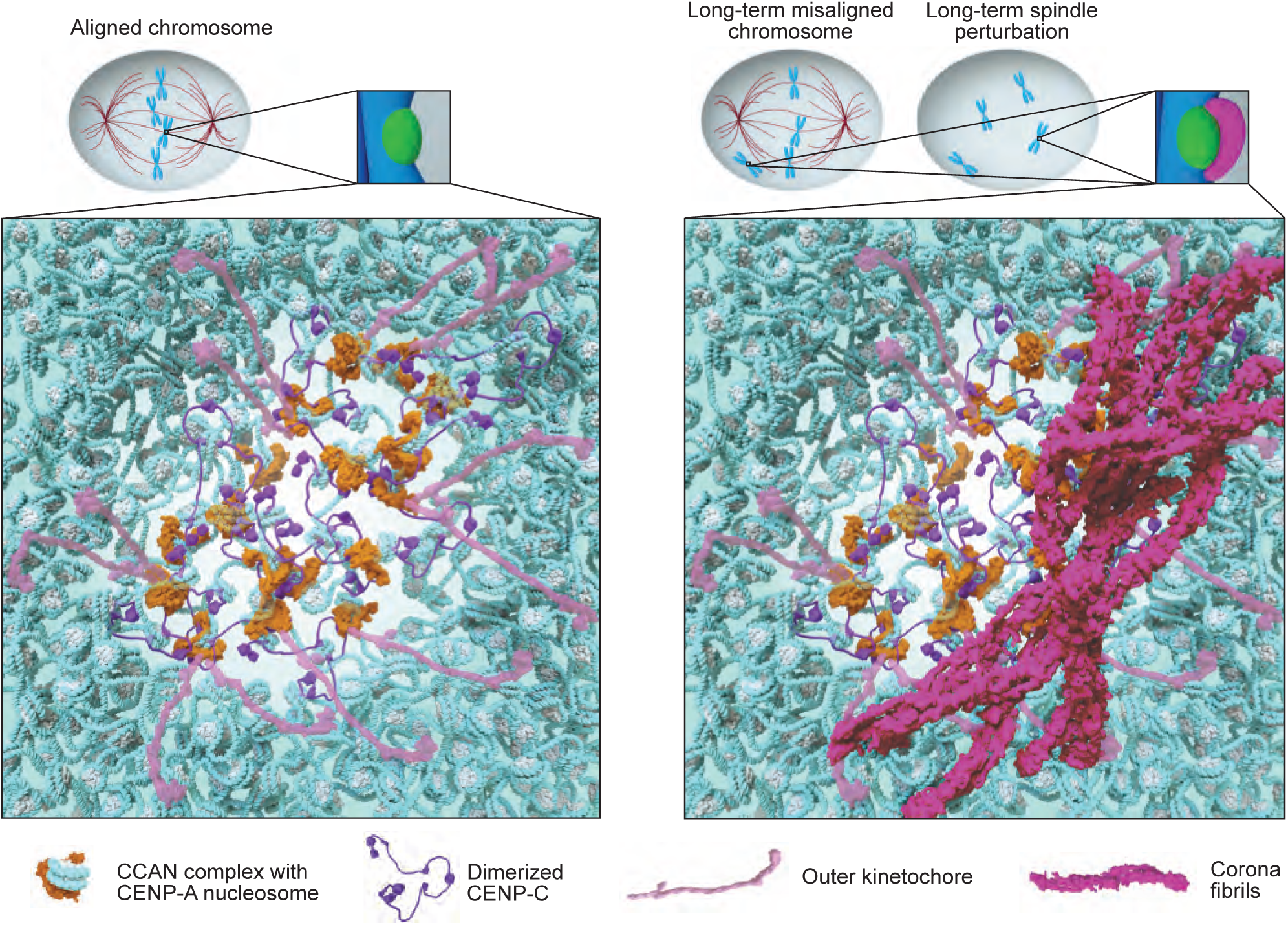
Models to describe the molecular architecture of human kinetochores. The inner kinetochore is clearly distinguished from bulk mitotic chromatin by CCAN complexes on the surface of the chromosome in a clearing devoid of dense chromatin. Early in mitosis or upon attachment to the mitotic spindle, the kinetochore lacks the corona. This arrangement is illustrated by the molecular model on the left. Note that we include illustration of the flexible mitotic couplers of the outer kinetochore, even though they are not readily distinguishable in our tomograms. If a chromosome does not readily attach and becomes trapped in a position where rapid attachment does not occur (or upon long-term spindle perturbation), the corona grows as part of a mitotic rescue mechanism. This arrangement is illustrated in the molecular model on the right. See text for details.

When a chromosome experiences prolonged detachment from the mitotic spindle, often occurring when a chromosome is not immediately attached to the spindle early in prometaphase and is partitioned in the cell entirely outside of the mitotic spindle (diagram in Fig. 8, right panel), the kinetochore is augmented by expansion of the fibrous corona (Fig. 8, right panel). The structural building blocks of the corona as visualized in our tomograms are likely RZZ-Spindly fibrils which form an extended meshwork. While there is regularity to the fibrous structure and a colinear arrangement of fibrils, the length of the individual fibrils is diverse, even within an individual kinetochore. Our tomograms reveal many unforeseen details of the organization of the fibrous corona, including how the fibrils interweave to form a crescent-shaped meshwork which arcs around the inner kinetochore forming a robust scaffold for binding of other corona components. There are strong biochemical and cell biological data that the RZZ-Spindly fibrous meshwork assembles high levels of components that capture spindle microtubules (e.g., CENP-E) and spindle assembly checkpoint activation (e.g., Mad2) (Kops and Gassmann, 2020). The crescent-like organization of these fibrils and their orientation tangential to the underlying chromatin and inner kinetochore as visualized in our tomograms contrasts with the radially oriented fibrous densities originally dubbed the fibrous corona from traditional EM images (Jokelainen, 1967). Our results suggest that these radially-oriented densities are instead the other components of the fibrous corona which dock on the RZZ-Spindly scaffold, observations which agree with recent models for corona organization (Kops and Gassmann, 2020). Thus, the tangentially-oriented RZZ-Spindly fibrils are more consistent with the “outer kinetochore plate” seen by traditional EM, which many decades ago was shown to expand with the corona and be composed of 10-20 nm fibrous domains oriented parallel/tangential to the underlying chromatin (Brinkley and Stubblefield, 1966; McEwen et al., 1993). The narrow, elongated complexes of the outer kinetochore, then, would connect these RZZ-Spindly fibrils to the inner kinetochore, consistent with the less-structured, electron-translucent region between the inner and outer kinetochore plates seen by traditional EM.

The three-dimensional visualization of the kinetochore we present here complements recent one-dimensional DNA maps of the chromatin landscape of the centromere locus (Altemose et al., 2022b; a; Dubocanin et al., 2023; Gershman et al., 2022; Logsdon et al., 2021; Miga et al., 2020). Areas of CENP-A occupancy coincide with regions of reduced CpG methylation termed centromere dip regions (Altemose et al., 2022a; b; Gershman et al., 2022; Logsdon et al., 2021; Miga et al., 2020), and Fiber-Seq of centromeres revealed these same regions to be highly physically accessible, forming the most accessible chromatin domain in the human genome (Dubocanin et al., 2023). These DNA sequencing-based results align remarkably well with our findings where tomograms show kinetochore complexes clustered within a distinct clearing space composed of open chromatin. Sequencing maps of human centromeres have also shown that CENP-A occupancy is highly variable in both magnitude and distribution, not only among individuals but also among chromosomes within an individual (Altemose et al., 2022a; Gershman et al., 2022; Logsdon et al., 2024). This attribute is also reflected in our tomograms where we observe kinetochores of varied size and with varied distribution of kinetochore complexes. Specifically, the finding that CENP-A occupancy is often split between multiple regions within a single centromere (Altemose et al., 2022a; Dubocanin et al., 2023; Gershman et al., 2022; Logsdon et al., 2024; Sacristan et al., 2022) is consistent with our own observation that some kinetochores are comprised of a single chromatin clearing whereas others are comprised of 2-3 clearings. Each of the clearings likely corresponds to a region on the linear DNA sequence with a high local density of CENP-A nucleosomes. How such apparent flexibility in the location of kinetochore-forming centromeric chromatin on the linear DNA could coincide with the faithful segregation of every chromosome during every cell division has remained a largely unanswered question. We conclude that a high local density of CENP-A nucleosomes leads to the formation of a higher-order structure capable of differentiating itself from the rest of the chromosome. Too sparse spacing of CENP-A nucleosomes on the linear DNA would prevent the local clustering of inner kinetochore complexes and minimize the ability to form distinct chromatin clearings. With sufficient density, a robust inner kinetochore is formed, leading to the formation of a structure that readily captures spindle microtubules and signals when they are not attached.

Visualization of macromolecular assemblies *in situ* has long provided the context to understand biochemical function, and the architecture of the kinetochore on mitotic chromosomes that we describe advances models to explain how chromosomes are faithfully inherited at cell division. We lastly note that our approach necessitated the development of a procedure to robustly target a chromatin domain that represents ∼0.03% of a typical chromosome. Thus, we anticipate that our approach could be adapted to targeting and imaging chromatin domains at other chromosome locations that are involved in other functional processes, such as those related to genome stability or gene expression.

## Acknowledgments

We thank E. Grishchuk (UPenn), M. Lampson (UPenn), and L. Jansen (Oxford) for comments on the manuscript. We thank our UPenn colleagues P. Allu for assistance with cell line generation, S. Mageswaran and B. Creekmore for assistance with cryo-ET data collection and analysis, and L. Guo for reagents. We thank S. Steimle of the Beckman Center for Cryo-Electron Microscopy and the Electron Microscopy Resource Lab Core Facility, RRID:SCR_022375, at UPenn for technical assistance. We thank D. Cleveland (UCSD), D. Fachinetti (Institut Curie), and I. Cheeseman (MIT) for reagents, and J. Iwasa (Utah) at the Animation Lab for the illustrations in Figure 8. This work was supported by NIH grants CA261198 (K.K.), GM134020 (Y.-W.C.), and GM130302 (B.E.B), a David and Lucile Packard Fellowship for Science and Engineering (2019-69645; Y.-W.C.), a Burroughs Wellcome Fund Investigators in the Pathogenesis of Infectious Disease Program award (1022785; Y.-W.C.), and a Pennsylvania Department of Health FY19 Health Research Formula Fund award (Y.-W.C.).

## Declaration of interests

The authors declare no competing interests.

## Author Contributions

All authors conceived the study, designed the experiments, analyzed the data, and wrote the manuscript. K.K. performed the experiments.

## Data and Materials Availability

Representative tomograms showing the human kinetochore under different conditions (Fig. 1D, Fig. 4D, Fig. 5D, Fig. 7B) will be made available in the Electron Microscopy Data Bank (EMDB) upon publication, under accession codes EMD-XXXXX (Fig. 1D), EMD-XXXXX (Fig. 4D, right panel), EMD-XXXXX (Fig. 5D), and EMD-XXXXX (Fig. 7D).

## Materials and Methods

### Generation of DLD-1 TIR1 CENP-A-rsEGFP2 cell line

The repair template for generation of the DLD-1 TIR1 CENP-A-rsEGFP2 cell line was constructed using the NEBuilder HIFI DNA Assembly Master Mix. Intronless CENP-A and the CENP-A 5’ UTR and 3’ UTR regions in pUC19 backbone were used from a previous EGFP-AID-CENP-A construct (Fachinetti et al., 2017). rsEGFP2 (Grotjohann et al., 2012) was ordered as a gBlock gene fragment (IDT). sgRNAs designed to target the 5ʹ and 3ʹ ends of the endogenous CENP-A gene were used in plasmids consisting of annealed oligos ligated into pX330, which contains Cas9(Ran et al., 2013). For the 5ʹ sgRNA, the following oligos were annealed: 5ʹ-CACCGgtgtcatgggcccgcgccgc -3ʹ and 5ʹ-AAACgcggcgcgggcccatgacacC -3ʹ. For the 3ʹ sgRNA, the following oligos were annealed: 5ʹ-CACCGcaactggcccggaggatccg-3ʹ and 5ʹ-AAACcggatcctccgggccagttgC-3ʹ. All plasmids were verified by sequencing. The CENP-A-rsEGFP2 repair template and sgRNA/Cas9 plasmids were co-transfected in a 9:1 ratio into DLD-1 TIR1 cells (Holland et al., 2012), a kind gift from D. Cleveland (UCSD), using Lipofectamine 2000 (Invitrogen). Transfected cells were screened by FACS for green fluorescence and sorted into single cells to generate monoclonal cell lines. A monoclonal line with biallelic CENP-A-rsEGFP2 was used for further work.

### Isolation of mitotic chromosomes

DLD-1 CENP-A-rsEGFP2 cells and DLD-1 CENP-C-AID-eYFP cells (Fachinetti et al., 2015) (a kind gift from D. Cleveland (UCSD) and D. Fachinetti (Institut Curie)) were cultured in Dulbecco’s Modified Eagle Medium (DMEM) supplemented with 10% fetal bovine serum, 100 U/ml penicillin, and 100 μg/ml streptomycin, maintained at 37° C and 5% CO_2_. Cells were arrested in mitosis by treatment with 50 μM S-trityl-L-cysteine (STLC) for 16 hours. 100 ng/mL nocodazole was added for the final 30 minutes of STLC treatment to depolymerize spindle microtubules. Mitotic cells were selectively harvested by mitotic blow-off, pelleted (600 × g, 5 min), and then allowed to swell for 10 min at room temperature in a solution of 5 mM PIPES pH 7.2, 10 mM NaCl, 5 mM MgCl_2_, 0.5 mM EGTA, and 2 mM EDTA. Cells were collected by centrifugation (600 × g, 5 min) and then resuspended in lysis buffer (5 mM PIPES pH 7.2, 10 mM NaCl, 5 mM MgCl_2_, 0.5 mM EGTA, 2 mM EDTA, 0.5 mM spermine, 1 mM spermidine, 10 mM NaF, 1 mM sodium orthovanadate, 1 mM PMSF, 0.0025 mg/mL leupeptin, 0.0025 mg/mL pepstatin, and 0.1% w/v digitonin) and lysed by 20 gentle strokes of a loose-fitting pestle. A cut pipet tip was used at this and all following steps to avoid chromosome shearing. The lysate was cleared by centrifugation (900 × g, 2 min), the concentration of NaCl increased to 100 mM, and the cleared lysate loaded onto a discontinuous sucrose gradient with 15%, 50%, and 80% (w/v) sucrose layers prepared in a buffer identical to the lysis buffer except 100 mM NaCl. The sucrose gradient was centrifuged at 5,000 × g for 15 min. Isolated chromosomes were removed from the 50%-80% interface, washed twice to remove sucrose (2,900 × g, 15 min), and then resuspended in 30 μL of the same buffer except without digitonin. Finally, the solution of isolated chromosomes was dialyzed into final chromosome buffer (5 mM PIPES pH 7.2, 50 mM NaCl, and 0.01% NP40) for 4 hours at room temperature. To image kinetochores under more-condensed chromatin conditions, chromosomes were isolated as above but dialyzed into a final buffer of: 5 mM PIPES pH 7.2, 50 mM NaCl, 0.25 mM MgCl_2_ and 0.01% NP40. For the CENP-C degradation experiment (CENP-C-AID-eYFP cell line (Fachinetti et al., 2015), +IAA condition), indole-3-acetic acid (IAA)(Sigma) was added at a final concentration of 500 μM for the final 1 hour of the mitotic arrest treatment to induce the degradation of CENP-C-AID-eYFP. Mouse anti-CENP-A monoclonal antibody (Enzo ADI-KAM-CC006-E; 1:1000) was added to the cleared lysate prior to running the sucrose gradient, and then cy5-conjugated Donkey anti-Mouse IgG (Jackson ImmunoResearch Laboratories #715-175-151; 1:200) was added to chromosomes after extraction from the sucrose gradient with unbound antibody removed by washing. For consistency, this antibody labeling was also used in the -IAA condition in which CENP-C-eYFP fluorescence is retained. For expansion of the fibrous corona, mitotic chromosomes were isolated from CENP-A-rsEGFP2 cells as described above except that the cells were arrested in mitosis using 100 ng/mL nocodazole alone for 16 hours. DLD-1 CENP-N-eGFP-AID cells (a kind gift from I. Cheeseman (MIT)) were cultured as described above for the other DLD-1 cell lines. For the CENP-N degradation experiment, cells were synchronized by single 2 mM thymidine block for 16 hours, released in 24 μM deoxycytidine for 5 hours, then arrested in mitosis with 50 μM S-trityl-L-cysteine (STLC) for 17 hours. 100 ng/mL nocodazole was added for the final 30 minutes of STLC treatment to depolymerize spindle microtubules. In the CENP-N degradation condition (+IAA condition), indole-3-acetic acid (IAA)(Sigma) was added at a final concentration of 500 μM for the final 12 hours of the mitotic arrest treatment to induce the degradation of CENP-N-eGFP-AID. Rabbit anti-CENP-C polyclonal antibody(Bassett et al., 2010) (1 μg/mL) was added to the cleared lysate prior to running the sucrose gradient, and then cy3-conjugated Goat anti-Rabbit IgG (Jackson ImmunoResearch Laboratories #111-165-144; 1:200) was added to chromosomes after extraction from the sucrose gradient with unbound antibody removed by washing. For consistency, this antibody labeling was also used in the -IAA condition in which CENP-N-eGFP fluorescence is retained.

### Cryo-ET grid preparation

A small portion of the isolated chromosomes was stained with DAPI and imaged in solution on a glass bottom dish with an inverted fluorescence microscope (Leica DMi8 or Leica DMI6000B, both with Leica DFC9000GT camera) to assess yield and check for retained chromosome morphology and kinetochore fluorescence. The remaining chromosome solution was then mixed with 200 nm diameter Tetraspeck beads (ThermoFisher Scientific) for CLEM and 10 nm colloidal gold fiducials (Ted Pella) for tilt-series alignment in tomogram reconstruction. 3 μL of chromosome solution was applied onto Quantifoil 200 mesh gold or copper R 2/2 extra thick carbon London Finder EM grids (Electron Microscopy Sciences). Excess liquid was blotted from the back (non-sample) side of the grid using Whatman filter paper #1. The grids were then plunge frozen into a liquid ethane/propane mixture using an EM GP2 plunge freezer (Leica Microsystems). Frozen grids were loaded into autogrid c-clip rings (ThermoFisher Scientific) and stored in liquid N_2_ for all subsequent steps.

### Correlative light and electron microscopy (CLEM)

EM grids were imaged by cryo-confocal microscopy on the Zeiss LSM900 Airyscan 2 with Quorum PP3010Z cryo-stage. Z-stacks were acquired on grid squares containing isolated chromosomes and maximum intensity projections generated. For the CENP-A-rsEGFP2 cell line, the equilibrium fluorescence of rsEGFP2 was used (without photoswitching) since this proved sufficient in brightness and localization accuracy for CLEM. Next, grids were transferred to a Thermo Fisher Titan Krios G3i 300 keV field emission cryo-transmission electron microscope with a K3 direct electron detector (Gatan Inc.). Imaging was performed using SerialEM (Mastronarde, 2005). Projection images of areas containing chromosomes were collected at 1,550x magnification with -50 μm defocus. The locations of 200 nm diameter Tetraspeck beads were marked in these images as well as in cryo-confocal images of the same regions, and then registration of the two coordinate systems was carried out in SerialEM (Mastronarde, 2005). CENP-A fluorescent foci were marked in confocal images and the corresponding positions in electron microscopy images were then targeted for cryo-ET data collection. This correlation was iteratively refined through a stepwise increase in magnification (correcting for small offsets between magnifications) up to a final magnification of 42,000x. After data acquisition, this correlation in SerialEM was revisited and refined to correct any small errors during initial on-the-fly correlation and data collection. The location of the center of the collected tilt-series relative to the targeted point was then used to generate overlays of cryo-confocal images with final tomogram slices.

### Cryo-ET data collection

Cryo-electron tomography (cryo-ET) was performed on a Thermo Fisher Titan Krios G3i 300 keV field emission cryo-transmission electron microscope with a K3 direct electron detector (Gatan Inc.), Volta phase plate (Danev et al., 2014), and an energy filter (slit width 20 eV at 42,000x, Gatan Inc.). Imaging was performed using SerialEM (Mastronarde, 2005). Target regions were initially assessed at lower magnifications and CLEM performed as described above. Cryo-ET data collection was performed at a nominal magnification of 42,000x (corresponding to a pixel size of 2.138 Å). Tilt-series were collected in a dose-symmetric scheme with a range of - 60° to +60° in 2° increments using the Volta phase plate and a defocus of -0.5 μm (Danev et al., 2017). For the more-condensed chromatin condition, the Volta Phase plate was not used, and the defocus was -6 μm. Tilt-series were collected with a cumulative dose of around 130 e-/Å^2^. Data on the two conditions using the CENP-C-AID-eYFP cell line (+/- IAA treatment) were collected sequentially during a single session on the Titan Krios, as were the two conditions using the CENP-N-eGFP-AID cell line (+/- IAA treatment). All other datasets were collected in individual sessions.

### Cryo-ET data processing and subtomogram averaging

Tilt-series were aligned using 10 nm colloidal gold as fiducial markers and reconstructed into tomograms using the IMOD software package (Kremer et al., 1996). Tomogram reconstruction was carried out by weighted-back projection after binning (bin4, voxel size 8.552 Å) using 10 iterations of the simultaneous iterative reconstruction technique (SIRT)-like filter. Tomograms that failed to reconstruct, showed excessively low contrast, or contained evidence of DNA shearing were excluded from analysis. The number of usable tomograms in each dataset are: 20 tomograms in the original CENP-A-rsEGFP2 dataset (Fig. 1-3,6), 20 tomograms for the CENP-C-AID-eYFP -IAA dataset (Fig. 4), 27 tomograms for CENP-C-AID-eYFP +IAA dataset (Fig. 4), 19 tomograms for CENP-N-eGFP-AID -IAA dataset (Fig. 5), 24 tomograms for CENP-N-eGFP-AID +IAA dataset (Fig. 5), and 21 tomograms for the CENP-A-rsEGFP2 dataset with expanded fibrous corona (Fig. 7). For subtomogram averaging of inner kinetochore (CCAN) complexes, model points were placed in the center of individual kinetochore complex densities identified by manual inspection, avoiding larger multimers or other nearby densities (corona fibrils, 10 nm fiducial beads, etc.). This yielded 141 particles from 9 tomograms. A more selective round of manual picking focused on the most clear and distinct inner kinetochore particles; this specifically involved exclusion of particles very close to other kinetochore complexes or (most commonly) very close to other nucleosomes and yielded 29 particles. In both cases, subtomogram averaging was carried out in PEET (Heumann et al., 2011; Nicastro et al., 2006) on bin4 tomograms (voxel size 8.552 Å) with a box size of 46 × 46 × 46 voxels using multiple iterations of rotational and translational searches. For subtomogram averaging of canonical nucleosomes, 190 particles were manually picked in regions away from kinetochore using IMOD. Each particle was manually pre-oriented (using the pattern of DNA gyres to determine nucleosome orientation) prior to subtomogram averaging and an initial average generated from pre-oriented particles was used as the initial reference for subtomogram averaging carried out as described above except with a box size of 20 × 20 × 20 voxels (voxel size 8.552 Å). The final positions and orientations of nucleosome particles were then used to generate initial positions for further subtomogram averaging from bin2 tomograms (voxel size 4.276 Å). The final positions and orientations of nucleosome particles from PEET at bin2 were then transferred into RELION 2 (Bharat and Scheres, 2016) for final 3D refinement and gold-standard resolution estimation by Fourier shell correlation. Note that particle identification and automated particle picking were attempted using 3D template matching methods for both inner kinetochore particles and canonical nucleosomes. However, we found this approach had unacceptably high false positive and false negative rates. We therefore concluded that, within the limitations of currently available software, 3D template matching is not suitable for unbiased identification of particles in the complex and crowded chromatin environment.

### Figure generation and modeling

Tomograms were oriented in 3D using IMOD’s Slicer window to capture the desired slice of a tomogram or individual particle. 10 layers of voxels (a thickness of 8.5 nm) were averaged around the section of interest to enhance contrast. Manual segmentation of kinetochore features was carried out in IMOD and displayed in IMOD or ChimeraX (Pettersen et al., 2021) for presentation. Trajectory of corona fibrils in tomograms was determined by visualization of the corona fibrils in tomograms and in lower magnification images acquired during CLEM, where tracts of fibrils bridging sister kinetochore are visible extending beyond the field of view of the final tomogram. The outlines of kinetochore chromatin clearings were segmented in IMOD for visualization of kinetochore shape. Subtomogram averages and PDB structures were manually oriented and displayed in ChimeraX for presentation. Line plots were generated in ImageJ/FIJI (Schindelin et al., 2012) after Gaussian filtering (σ = 2.0) of the tomogram slice and plotted with Prism 9.

### Quantification of kineotochore architecture

Analysis of 2D tomogram slices in ImageJ/FIJI (Schindelin et al., 2012) was used to quantitatively compare kinetochore loci to surrounding chromatin. Snapshots were acquired of single tomogram slices containing kinetochore loci at 1x magnification in IMOD (Kremer et al., 1996). In FIJI, each snapshot was cropped to 600 × 600 pixels^2^ surrounding the kinetochore region, filtered with a Gaussian blur (σ = 1.0), and then automatically thresholded using the “MaxEntropy” method to generate a binary image. A filter for small particles (50-250 pixels^2^) was applied to this binary image, and the region largely excluding such small particles (i.e., nucleosomes) was used to initially define the kinetochore region (chromatin clearing). The boundary of the kinetochore region was manually refined and then this boundary was used to define the kinetochore vs. outside-kinetochore regions of the image. For both regions, particles above 50 pixels^2^ in area were analyzed to calculate particles per unit area and determine particle area (nm^2^). The NND plugin was used to calculate nearest neighbor distance between the centroids of particles within the selected region. For each tomogram slice, the particles per 1000 nm^2^, average particle area (nm^2^), and average NND (nm) were calculated for both the kinetochore and outside-kinetochore regions. 5 non-sequential slices spaced through the kinetochore area of each of 10 tomograms were analyzed in this way (N=50). This analysis was carried out on 2D slices rather than 3D volumes because the missing wedge causes elongation of particles along the tomogram z-axis and so automated segmentation cannot distinguish particles which are in a similar x-y position but separated in z, leading to the creation of artifactual “large particles” that extend through much of the z-thickness of the tomogram. This phenomenon adds an unacceptable degree of noise to any attempts at 3D quantification. For all comparisons, a paired two-tailed t-test was used to compare kinetochore vs. outside-kinetochore regions. Each kinetochore region was paired to the outside-kinetochore region from the same tomogram slice. Descriptive statistics were also calculated. For quantification of the number of larger densities (complexes) per kinetochore, 10 representative tomograms were chosen from the partially decondensed chromatin and condensed chromatin datasets. The distinct larger densities within the entirety of the kinetochore clearing(s) were counted, with larger multimer densities being counted as a single particle. For the few tomograms in which part of the non-targeted sister kinetochore is visible within the field of view, only the targeted kinetochore was assessed. For assessment of kinetochore volume and sphericity, 5 representative tomograms were chosen from each dataset, excluding any kinetochores with more than one chromatin clearing or instances in which the kinetochore intersects with the edge of the tomogram. The boundary of the chromatin clearing was manually segmented across multiple XY tomogram slices and then interpolated in IMOD(Kremer et al., 1996). The volume and surface area of this segmentation were used to calculate sphericity. Unpaired t-tests were used to compare partially decondensed chromatin and condensed chromatin conditions for these data. For quantification of the volume of individual kinetochore complexes, complex densities were manually segmented, and their volume was assessed in IMOD. The number of clear nucleosomes (identified by density corresponding to DNA gyres with characteristic pattern of wrapping an octameric nucleosome) within each complex was then counted. The relationship between complex volume and nucleosome number was assessed by simple linear regression.

All statistical analysis was carried out in Prism 9. Data are presented as mean ± standard deviation (SD).

### Immunofluorescence

DLD-1 cells were fixed in 4% paraformaldehyde for 10 min at room temperature, quenched with 100 mM Tris (pH 7.5) for 5 min, and then permeabilized using PBS with 0.1% Triton X-100. Coverslips were blocked in PBS supplemented with 2% fetal bovine serum, 2% bovine serum albumin, and 0.1% Tween 20 before antibody incubations. The following antibodies were used, for CENP-A immunofluorescence: mouse anti-CENP-A mAb (Enzo ADI-KAM-CC006-E; 1:1000) and cy5-conjugated Donkey anti-Mouse IgG (Jackson ImmunoResearch Laboratories #715-175-151; 1:200), for CENP-C immunofluorescence: rabbit anti-CENP-C polyclonal antibody (Bassett et al., 2010) (1 μg/mL) and cy3-conjugated Goat anti-Rabbit IgG (Jackson ImmunoResearch Laboratories #111-165-144, 1:200), for CENP-T immunofluorescence: rabbit anti-CENP-T polyclonal antibody (Hori et al., 2008) (1 μg/mL), a kind gift from Iain Cheeseman (MIT), and cy3-conjugated Goat anti-Rabbit IgG (Jackson ImmunoResearch Laboratories #111-165-144; 1:200), and for CENP-E immunofluorescence: mouse anti-CENP-E mAb177 (Santa Cruz 47745; 1:100) and cy5-conjugated Donkey anti-Mouse IgG (Jackson ImmunoResearch Laboratories #715-175-151; 1:200). Coverslips were stained with DAPI before mounting with VectaShield medium (Vector Laboratories). Z-stacks were acquired of cells and projected as a single 2D image by maximal intensity projection for presentation. For images of purified chromosomes, antibody labeling was carried out during chromosome isolation as described above, the following antibodies were used: for CENP-C: rabbit anti-CENP-C polyclonal antibody (Bassett et al., 2010) (1 μg/mL) and cy3-conjugated Goat anti-Rabbit IgG (Jackson ImmunoResearch Laboratories #111-165-144; 1:200), for Ndc80/Hec1: mouse anti-Hec1 monoclonal antibody (Abcam ab3613, 1:1000) and cy5-conjugated Donkey anti-Mouse IgG (Jackson ImmunoResearch Laboratories #715-175-151; 1:200). Chromosomes in solution were imaged on coverslips without fixation or mounting medium. Images were captured on an inverted fluorescence microscope (Leica DMi8 or Leica DMI6000B, both with Leica DFC9000GT camera) and a 100x oil-immersion objective. Chromosomes labeled for CENP-C and Ndc80 were also imaged frozen onto EM grids for visualization and quantification of kinetochore fluorescence intensity. Quantification of fluorescence intensity relative to background was determined in ImageJ/FIJI (Schindelin et al., 2012) for each channel. Overlays of multiple channels were assembled using ImageJ/FIJI.

---

**Figure S1:**
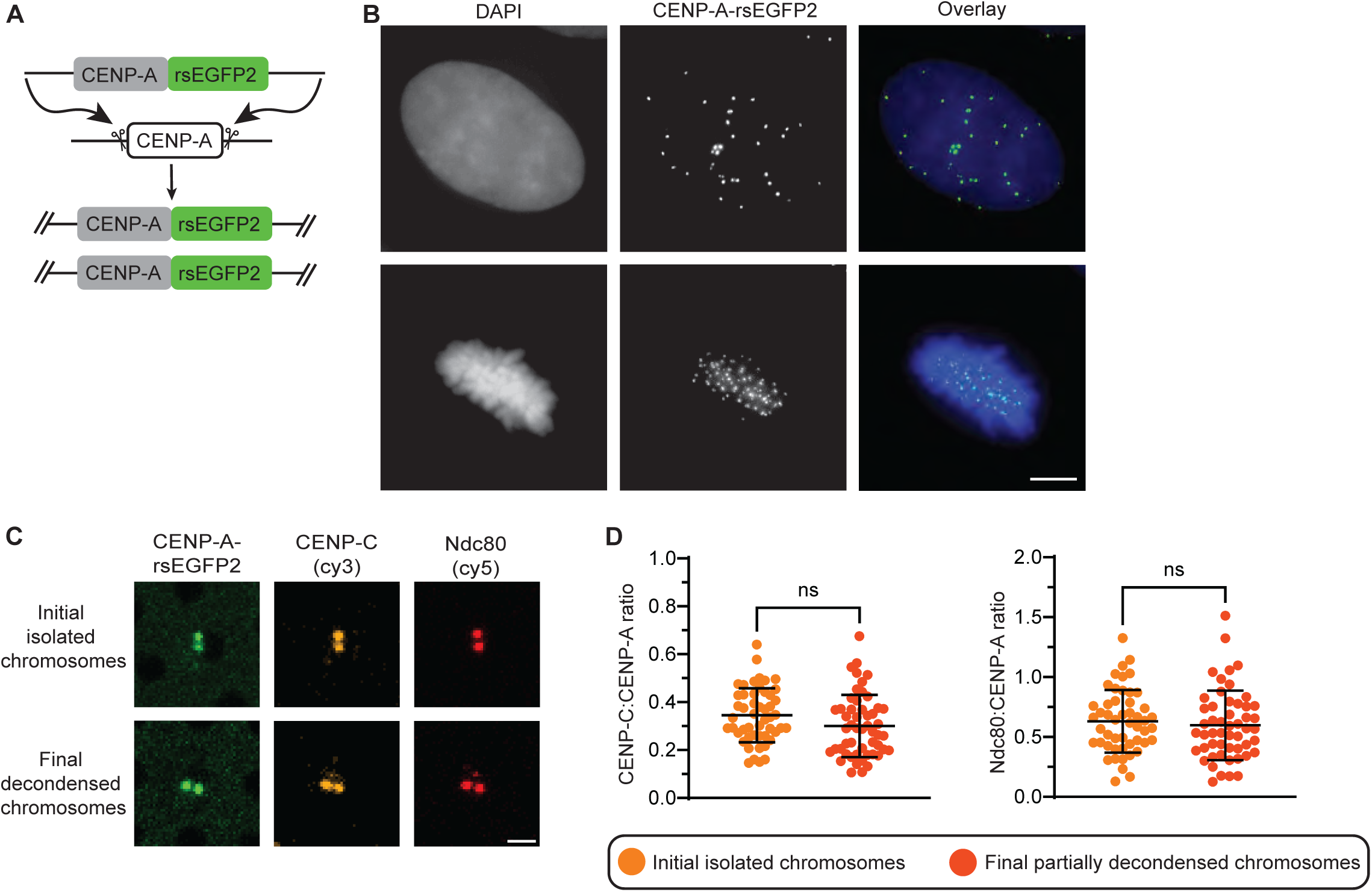
CENP-A-rsEGFP2 cells and isolated chromosomes. **A.** Diagram of the generation of DLD-1 CENP-A-rsEGFP2 cell line. **B.** Example fluorescence images of DLD-1 CENP-A-rsEGFP2 cell line, showing representative interphase (upper panels) and metaphase (lower panels) cells. Scale bars, 5 μm. **C.** Cryo-confocal images showing centromeric fluorescence from initial isolated chromosomes (fully condensed, immediately after isolation from the cell) and final, partially decondensed chromosomes (such as those used for cryo-ET data collection) from CENP-A-rsEGFP2 cell line (frozen on EM grids: in the absence of DAPI staining, chromosomes were identified by their shape in T-PMT channel). The chromosomes are labeled by immunofluorescence for CENP-C and Ndc80 (Hec1). Both CENP-C, an inner kinetochore component, and Ndc80, an outer kinetochore component, are retained on isolated chromosomes and co-localize with CENP-A. Scale bar, 2 μm. **D.** Quantitative comparison of CENP-C and Ndc80 fluorescence intensity on initial isolated chromosomes and final partially decondensed chromosomes as in (C) measured on chromosomes frozen on EM grids. Data is represented as ratio of CENP-C or Ndc80 fluorescence to CENP-A fluorescence at the same centromere. These data show that inner and outer kinetochore components are quantitatively retained at the kinetochore during isolated chromosome preparation and decondensation.

**Figure S2:**
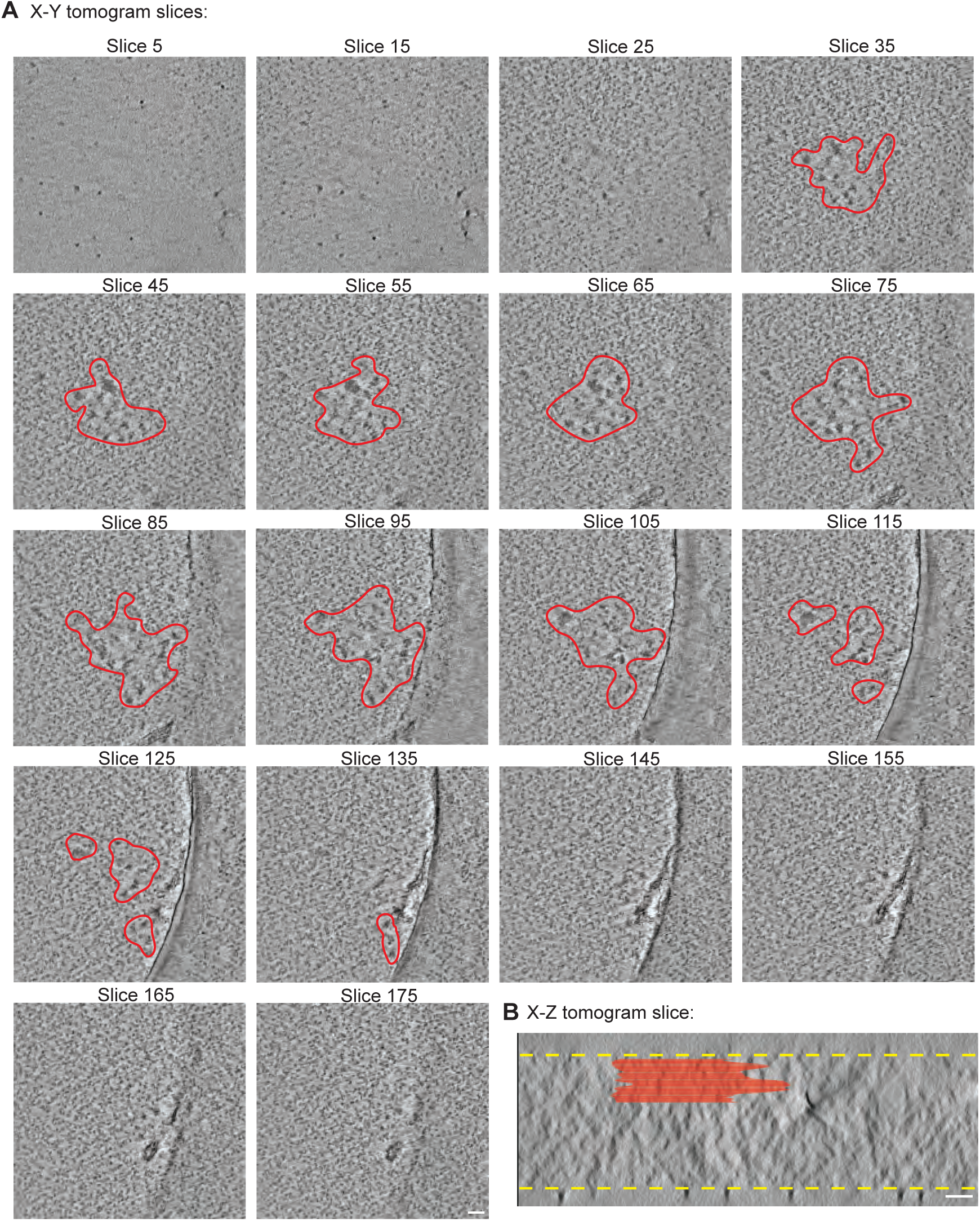
Kinetochores are located at the chromosome surface. **A.** A series of x-y slices from the tomogram shown in Figure 1D-E showing that the kinetochore is located at the surface of the chromosome. Slices are separated in the z-axis by about 8.5 nm and traverse from vitreous ice above the chromosome surface through the full kinetochore area and finish in a region of ordinary mitotic chromatin past the kinetochore. The chromatin clearing is outlined in red. For further details see Video S1. Note that the carbon film, the edge of a hole in the film, and torn carbon fragments are visible in slices 65-175. Scale bars, 50 nm. **B.** A side view (x-z plane) of the full tomographic volume from (A) overlaid with the segmentation of the kinetochore area in red as in (A). This shows that the kinetochore is located at the chromosome surface. The surfaces of the vitreous chromosome are shown with a yellow dashed line.

**Figure S3:**
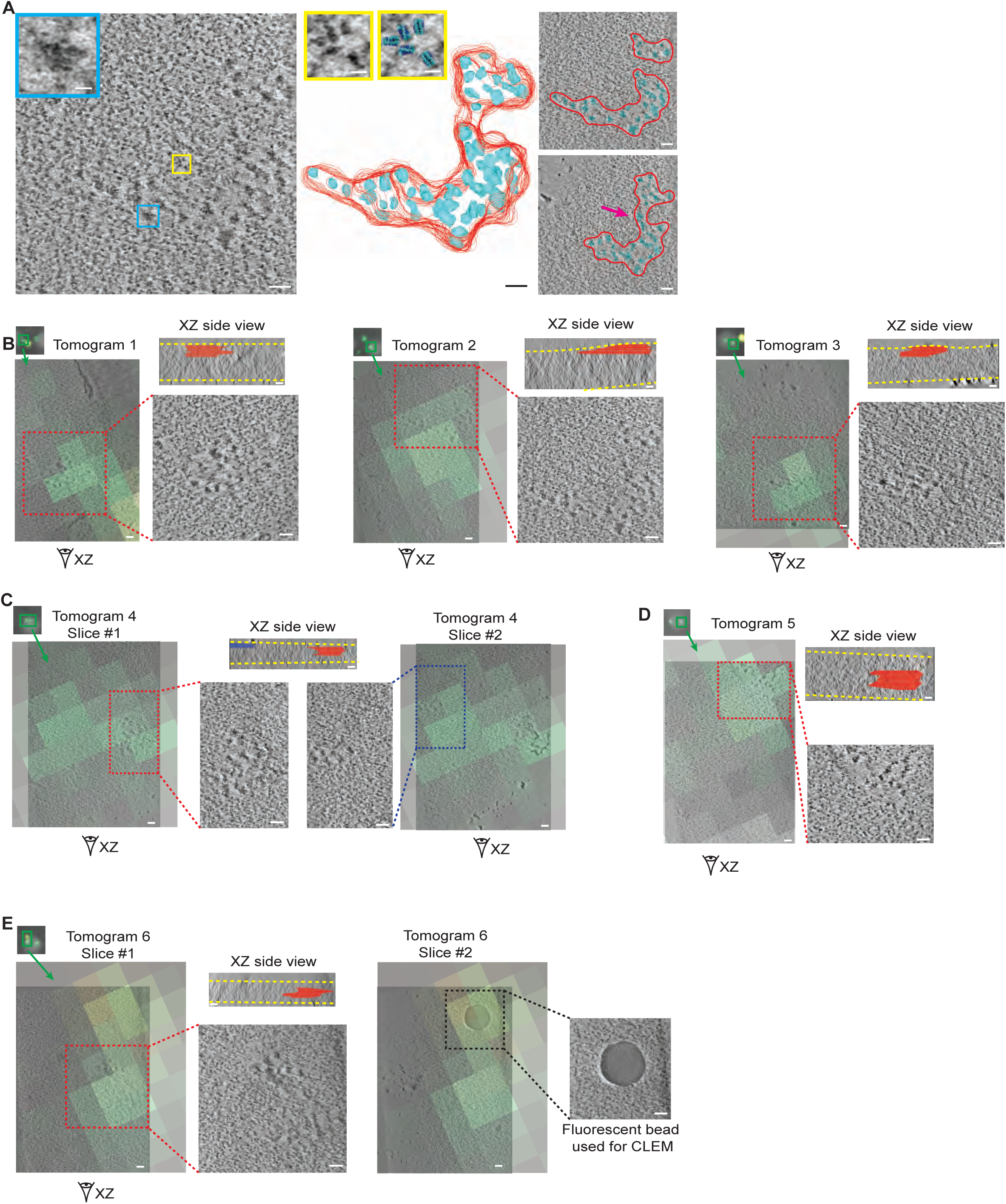
Tomograms targeted to CENP-A-rsEGFP2 fluorescence show consistent kinetochore architecture. **A.** An additional example tomogram showing a kinetochore annotated as in Figure 1D-E. On the left is a tomogram slice showing kinetochore complexes within chromatin clearings, in the center is a 3D model of the full locus, and to the right is overlay of the model with other slices of the same tomogram. This kinetochore is comprised of two chromatin clearings which coalesce at the surface of the chromosome (pink arrow). In the model, the boundaries of the chromatin clearings are traced in red and kinetochore complexes are traced in cyan. Cyan inset shows an example kinetochore complex. Yellow inset shows a region of chromatin outside the kinetochore with an overlay of nucleosome structures (PDB: 1KX3). Scale bars, 50 nm (insets 10 nm). **B-E.** Examples of kinetochores seen in tomograms targeted to CENP-A-rsEGFP2 fluorescence, demonstrating the accuracy of correlation and the consistent architecture of kinetochores. In each case the “double dot” fluorescence pattern of a mitotic chromosome is shown and the kinetochore that was imaged is highlighted with a green box. A slice of the resultant tomogram is overlaid with the fluorescent signal at that position. The location of the kinetochore within the tomogram is shown by a red or blue dashed box and expanded in the inset. In all cases, we find the distinct chromatin architecture of the kinetochore in very close proximity to the center of CENP-A fluorescence after CLEM. The 3D location of the kinetochore within each tomogram is demonstrated with a side view of the tomogram. In these side views, the kinetochore volume is outlined in red or blue and the edge of the vitrified chromatin is outlined in yellow. Scale bars, 50 nm. Several experiments testing immunogold labeling with various diameters of gold beads were attempted. Small beads (1.4 nm) conjugated with fluorescent moieties properly localized to kinetochores but these beads were too small to clearly identify in tomograms amongst DNA densities with similar molecular radii. Larger beads did not properly label the kinetochore, likely due to their inability to penetrate near the surface of chromosomes. **C.** Tomogram 4 captures parts of two sister kinetochores (the kinetochores of sister chromatids of a single chromosome) within the same tomogram. The targeted kinetochore is shown in red and the other sister kinetochore (only a fraction of which is contained within the tomogram field of view) is shown in blue. The sister kinetochores are located in separate chromatin clearings but share the characteristic architecture of the kinetochore. The relative positioning of the two sister kinetochores in 3D can be seen in the x-z side view of the tomogram. **D-E.** Tomograms 5 and 6 capture kinetochores in which the inter-kinetochore axis is parallel to the underlying EM grid and so the kinetochore is positioned in the middle of the tomogram z-thickness (rather than at a surface), as shown in the x-z side view of the tomograms. **E.** Tomogram 6 captures a kinetochore as well as one of the fluorescent beads used for CLEM, in separate slices of the tomogram. Fluorescent overlay highlights the accuracy of correlation between confocal and electron microscopy images. In slice #1, the CENP-A green fluorescence coincides with the kinetochore. In slice #2, the yellow fluorescence (representing the bead which fluoresces in multiple channels) is centered on the physical bead.

**Figure S4:**
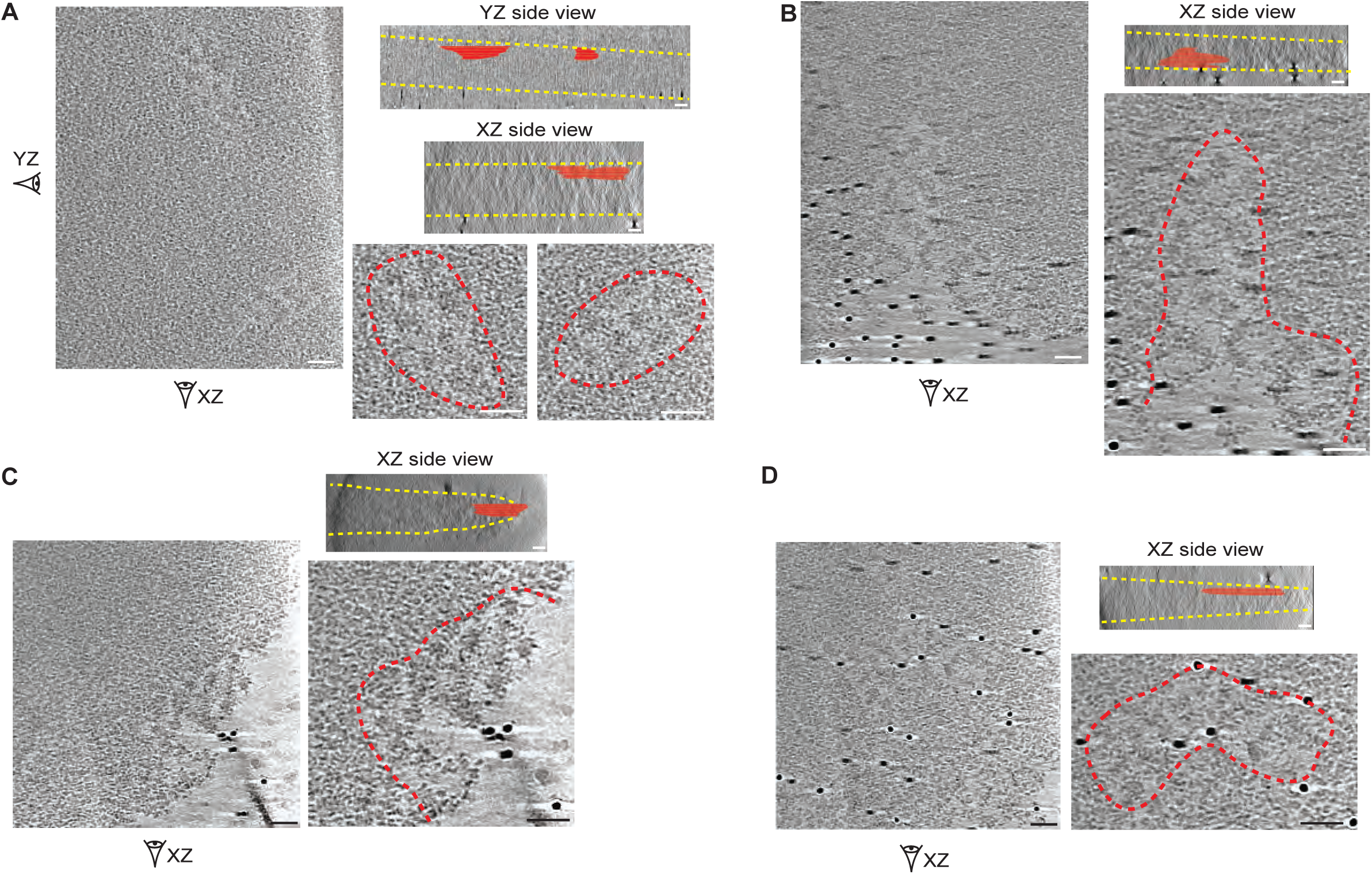
Kinetochore chromatin clearings are observed on heavily-condensed mitotic chromosomes, and some kinetochores contain more than one chromatin clearing. **A-D.** Example tomogram slices showing kinetochores from isolated chromosomes prepared under conditions of condensed chromatin (see Methods). The kinetochore areas are expanded in insets and outlined with red dashed lines. Tomogram slices are accompanied by side views of the tomogram in which the kinetochore volume is outlined in red and the edge of the vitrified chromatin is outlined in yellow. Under these conditions of more-compacted chromatin, little to no open space is visible between nucleosomes in surrounding chromatin. However, chromatin clearings are still evident with open space surrounding larger inner kinetochore complexes. This demonstrates that the chromatin clearings seen in Figure 1 are not only found when there is increased chromatin decondensation, as in that preparation. **A.** This single kinetochore is composed of two distinct chromatin clearings. B-D. The orientation of the kinetochore relative to the chromosome edge more easily appreciated under conditions of condensed chromatin (in which the chromosome is more rigid and thus more likely to lay with sister kinetochores parallel to the underlying carbon film). **B,C.** Two examples of kinetochores from chromosomes oriented with the inter-sister kinetochore axis roughly parallel to the underlying EM grid. In these cases, a true “side view” of the kinetochore at the chromatin surface is evident with the kinetochore complexes seen at the edge of the chromatin within an x-y tomogram slice. In tomograms with similar orientations under conditions of more decondensed chromatin, the chromosome edge can be obscured by blotting of surrounding chromatin around the kinetochore (Fig. S3D,E). **D.** An example kinetochore from a chromosome slightly oblique relative to the underlying EM grid. The kinetochore is seen proximal to the chromosome edge (lower right corner of tomogram) but due to the chromosome orientation it sits at the “top” surface of the chromatin in x-z side views and is surrounded by ordinary mitotic chromatin in x-y slices. As in (B,C), this effect is exaggerated under conditions of increased chromatin decondensation, which can obscure kinetochore orientation relative to the chromosome edge. Note that 10 nm gold fiducial beads are visible in panels B-D. Scale bars, 50 nm.

**Figure S5:**
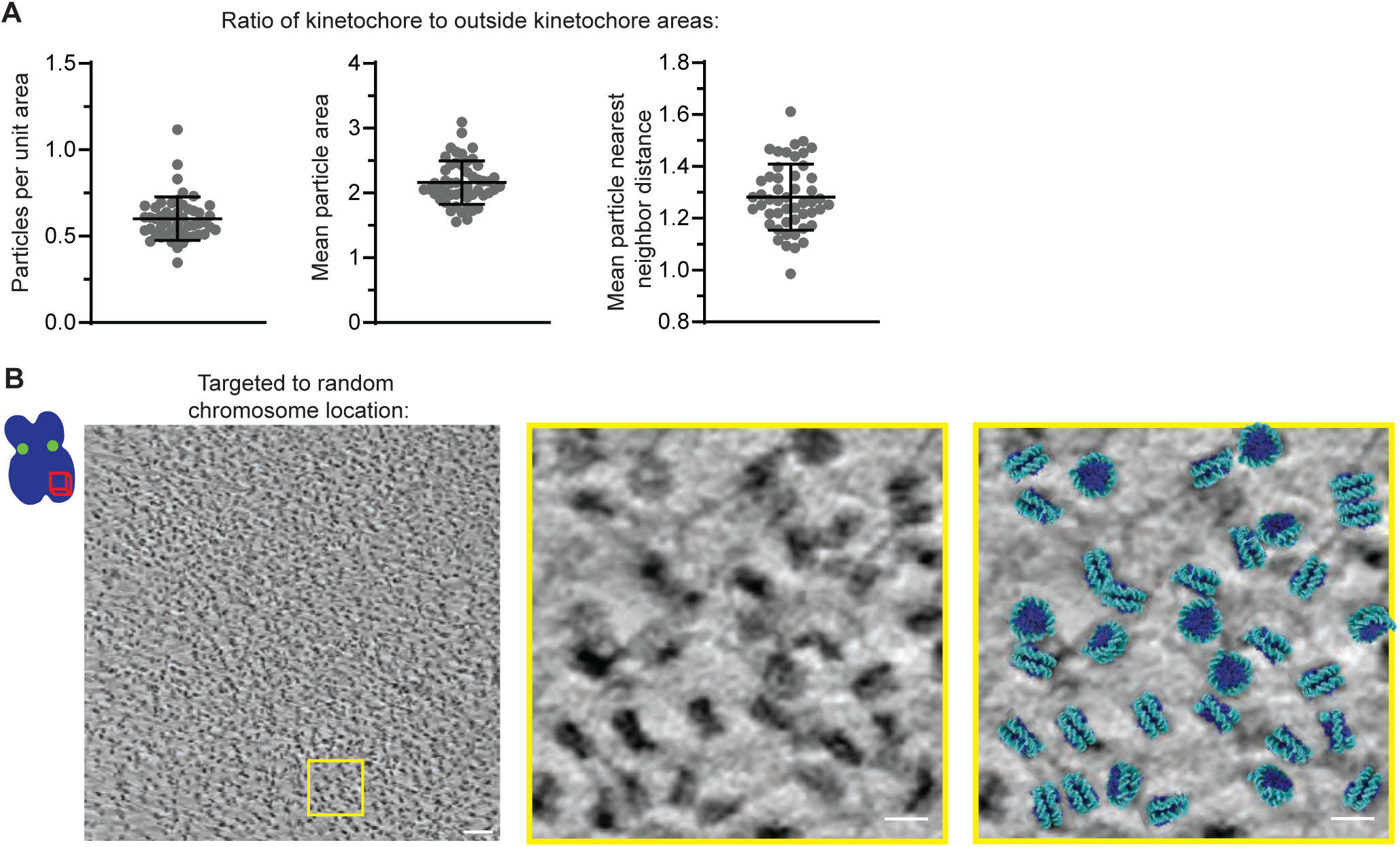
Comparing kinetochores to surrounding chromatin. **A.** Data from Figure 2C-E expressed as ratio values between kinetochore area and surrounding chromatin. Mean ± SD: Particles per 1000 nm^2^: 0.60 ± 0.13. Mean particle area (nm^2^): 2.16 ± 0.34. Mean particle NND (nm): 1.28 ± 0.13. **B.** Representative slice of a tomogram captured at a location on the chromosome arm away from CENP-A fluorescence. Inset area shows nucleosomes and DNA strands, paired with an overlay of the structure of an octameric nucleosome (PDB: 1KX3). This tomogram shows the architecture of mitotic chromatin at all chromosome locations away from CENP-A fluorescence imaged in this study and consists solely of packed canonical nucleosomes. Scale bar, 50 nm (insets 10 nm)

**Figure S6:**
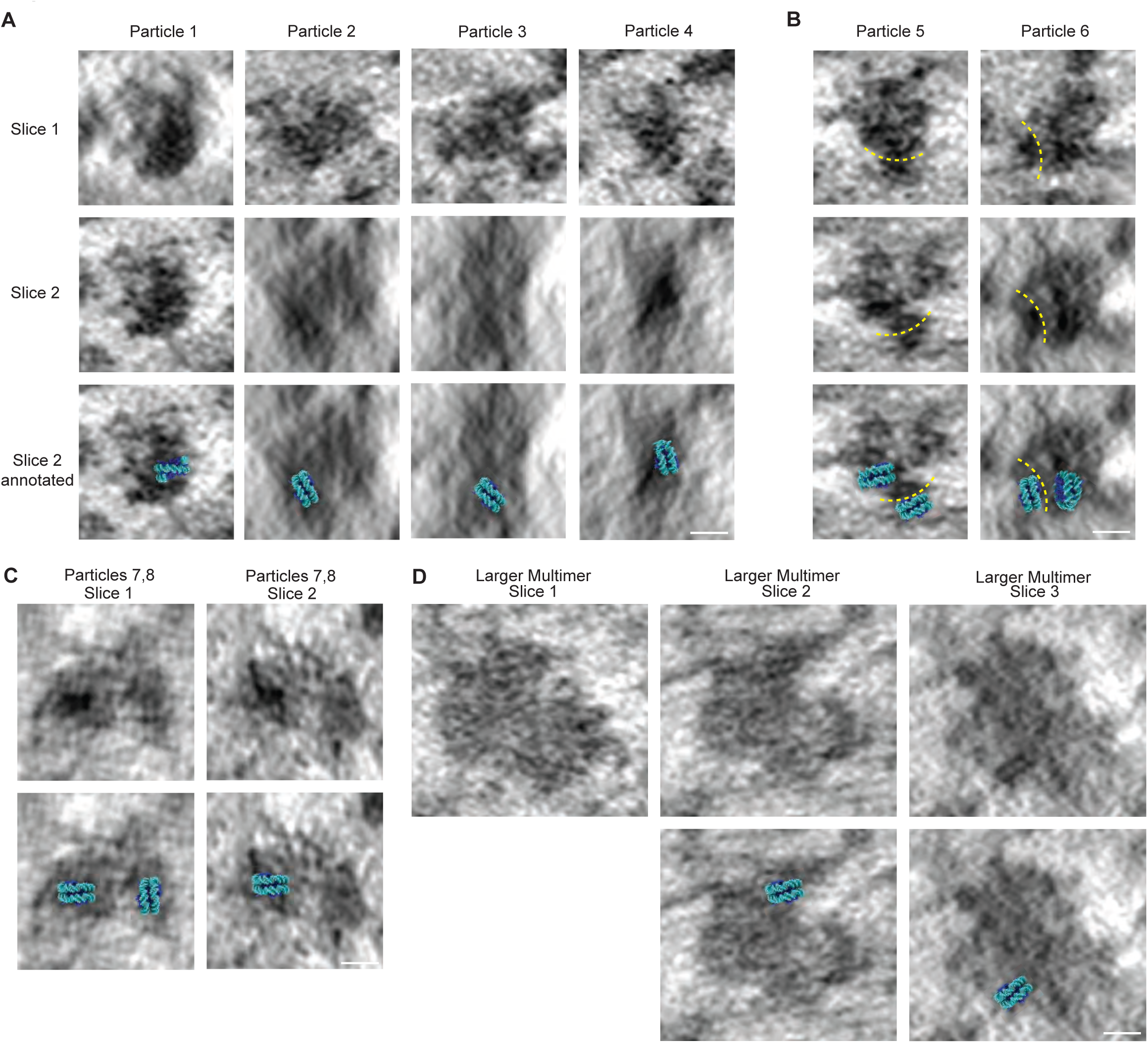
A nucleosome with two gyres of wrapped DNA is embedded in individual kinetochore complexes and multimers thereof. **A.** Further examples of individual inner kinetochore complexes with clear density corresponding to a nucleosome embedded within the complex, as in Figure 3C. For each particle, slice 2 shows the particle rotated to an angle which allows visualization of the DNA gyres wrapping the nucleosome particle. In all cases, two DNA gyres are visualized, consistent with an octameric nucleosome. Scale bars, 10 nm. **B.** Examples of individual kinetochore complexes with a nucleosome embedded within the complex and a second nucleosome closely associated with the complex. The second nucleosome likely represents the intervening nucleosome between two kinetochore complexes. Yellow dashed lines separate the density of the kinetochore complex from that of the second nearby nucleosome. Nearby nucleosomes such as these are a major contributor to the visual heterogeneity of inner kinetochore complexes in tomograms. **C,D.** Higher order packing of inner kinetochore complexes is observed in our tomograms which provides a high density of binding sites for outer kinetochore elements. **C.** Example tomogram slices showing two closely associated kinetochore complexes, both of which contain density corresponding to a nucleosome. **D.** Tomogram slices showing a multimer of kinetochore complexes (also shown in Fig. 3H). Multiple nucleosome densities are observed within this larger multimer, and two examples are shown here.

**Figure S7:**
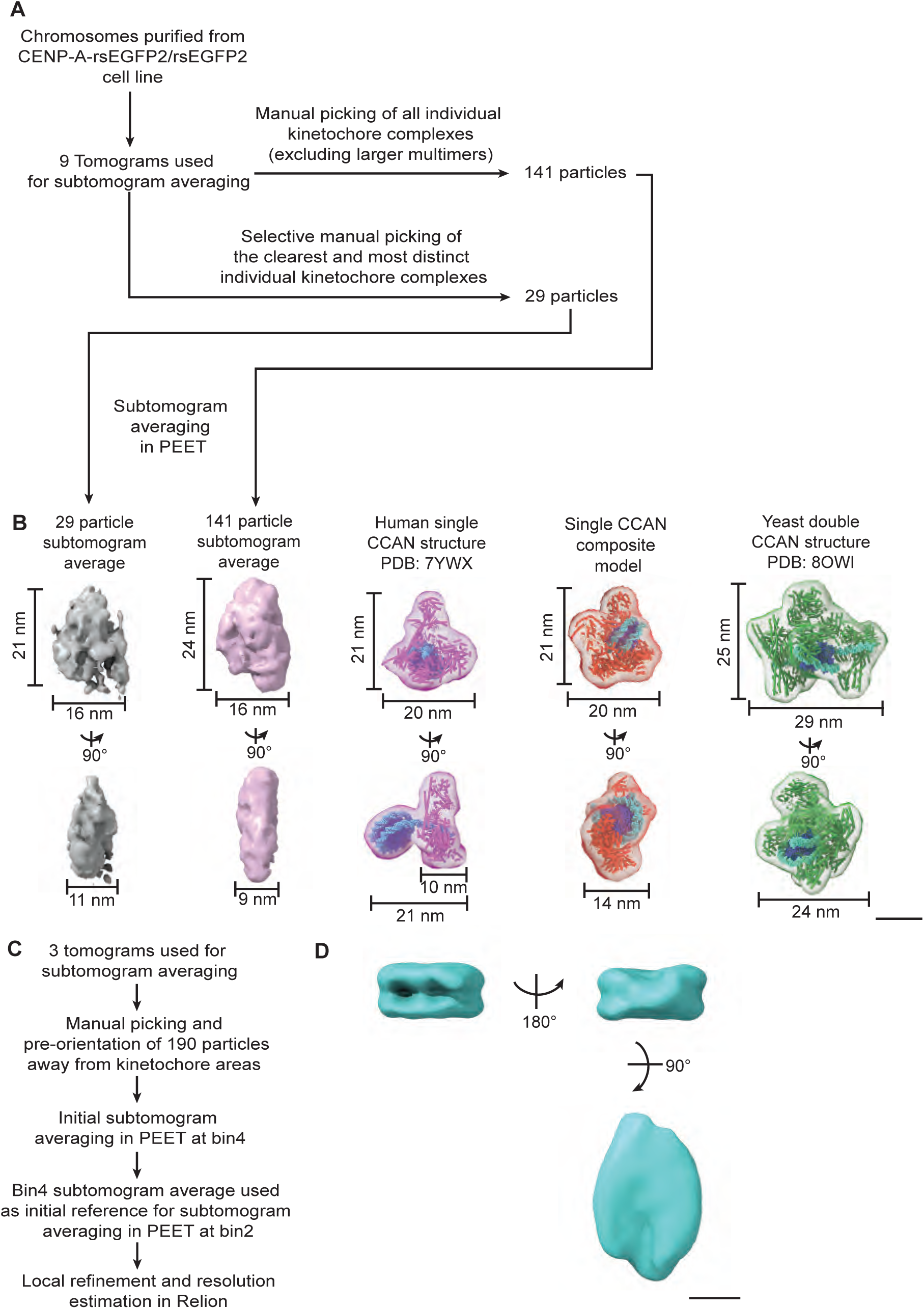
Subtomogram averaging of CCAN complexes and canonical nucleosomes. **A.** Schematic of subtomogram averaging workflow used for CCAN complexes. An initial round of manual particle picking identified 141 particles from 9 tomograms which represented individual kinetochore particles (i.e., avoiding multimers) and were suitable for subtomogram averaging (i.e., not near the tomogram edge, not near 10 nm gold fiducial beads). A second round of manual particle picking was more selective for inner kinetochore particles which were clear and distinct from their surroundings. This round of particle picking specifically avoided particles such as those in Figure S6B in which nearby nucleosomes are closely associated with the inner kinetochore complex, creating a seemingly-larger particle. **B.** Two low-resolution subtomogram averages were obtained using the different sets of kinetochore particles described in (a). Dimensions of both subtomogram averages are shown alongside the two reconstituted inner kinetochore structures from Figure 3F: human (PDB 7YWX)(Yatskevich et al., 2022) and yeast (*S. cerevisiae*)(PDB 8OW1)(Dendooven et al., 2023), as well as a yeast-human hybrid model of a single CCAN complex with embedded CENP-A nucleosome (Kixmoeller et al., 2020). Each structure or model is shown within a 50 Å resolution envelope. The subtomogram averages are most consistent in size and shape with a single copy of the CCAN. Scale bar, 10 nm. **C.** Schematic of subtomogram averaging workflow used for individual canonical nucleosomes picked from regions away from the kinetochore. Particles were manually pre-oriented using DNA gyres to determine particle orientation. **D.** Three views of the 24 Å resolution average obtained from 190 nucleosome particles. The rigid and consistent nature of nucleosome particles made them more easily averaged than inner kinetochore particles. Scale bar, 5 nm.

**Figure S8:**
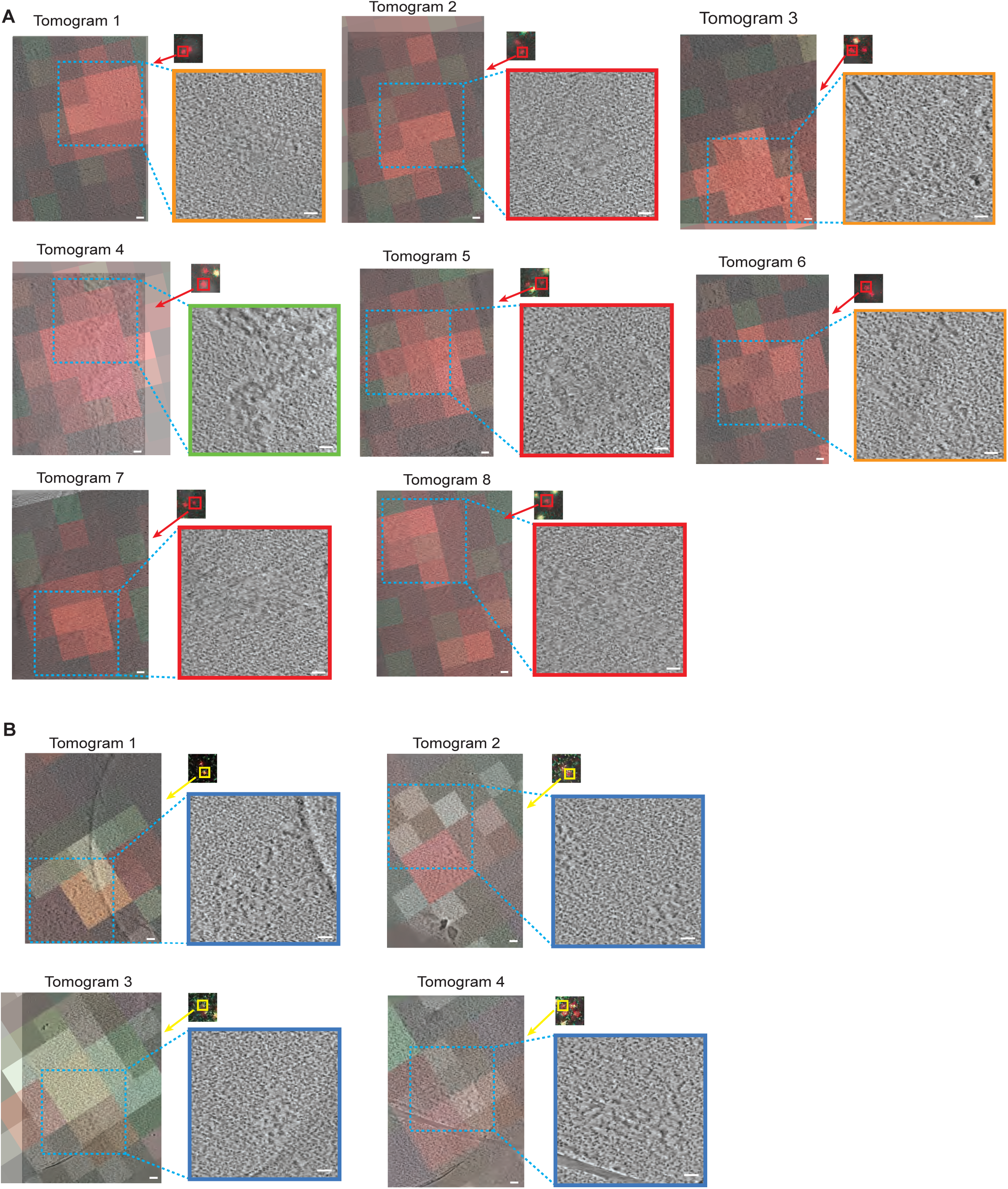
Further examples of kinetochores from CENP-C-AID-eYFP cells. **A.** Example tomograms from the CENP-C-AID-eYFP +IAA condition, in which CENP-C is degraded, showing that removal of CENP-C perturbs kinetochore architecture. In each case the “double dot” CENP-A fluorescence pattern (from antibody labeling) of a mitotic chromosome is shown and the kinetochore that was imaged is highlighted with a red box. A slice of the resultant tomogram is overlaid with the fluorescent signal at that position, demonstrating the accuracy of CLEM. In all cases, we identify the kinetochore at the area of CENP-A fluorescence. The location of the kinetochore locus within the tomogram is shown by the blue dashed box and expanded in the inset. The majority of kinetochores in this condition of CENP-C degradation show perturbation of kinetochore architecture. Each kinetochore inset is colored according to its morphology as defined in Figure 4E. Tomogram 8 shows an example of a kinetochore with total loss of distinct architecture in which the kinetochore is positioned over a hole in the EM grid carbon film. Scale bars, 50 nm. **B.** Kinetochore architecture is preserved in CENP-C-AID-eYFP cells without CENP-C degradation as shown in example tomograms from the CENP-C-AID-eYFP - IAA condition. In this condition CENP-C is not degraded, and as expected the architecture of the kinetochores matches that seen in our initial preparation from CENP-A-rsEGFP2 cells (Fig. 1,2, S2,S3). Tomogram slices are shown as in (A) except that fluorescent foci are yellow due to colocalization of CENP-A-cy5 and CENP-C-eYFP.

**Figure S9:**
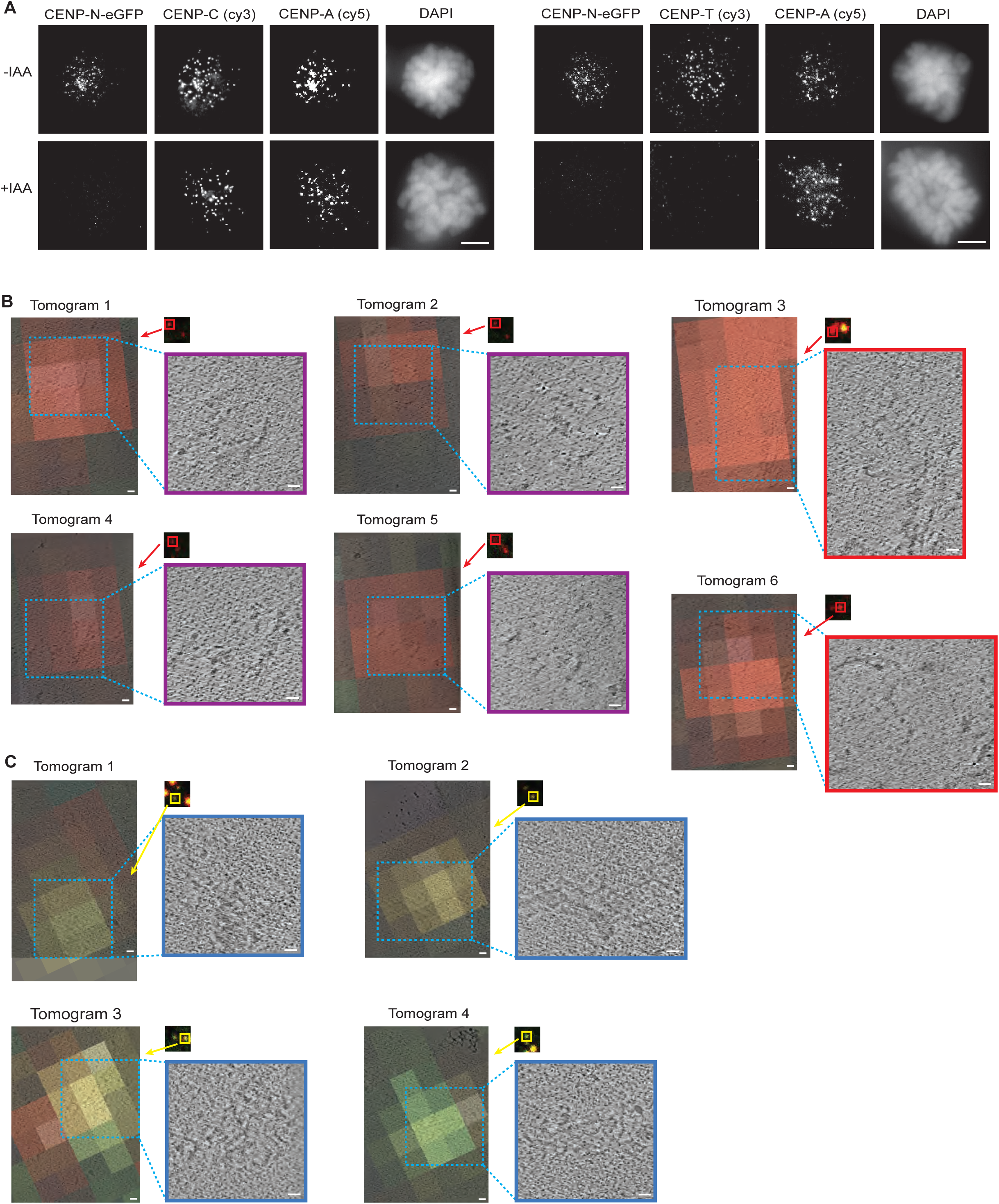
Further examples of kinetochores from CENP-N-eGFP-AID cells. **A.** Immunofluorescence images from CENP-N-eGFP-AID cells synchronized in mitosis, with and without the addition of IAA to degrade CENP-N (as shown in Fig. 5B) with immunolabeling of CENP-A and either CENP-C or CENP-T. A representative cell is shown for each condition. After mitotic degradation of CENP-N, CENP-C is retained at the centromere while CENP-T is lost. Scale bars, 5 μm **B.** Example tomograms from the CENP-N-eGFP-AID +IAA condition, in which CENP-N is degraded, showing that removal of CENP-N reduces the size of kinetochore complexes while leaving the chromatin clearing intact. In each case the “double dot” CENP-C fluorescence pattern (from antibody labeling) of a mitotic chromosome is shown and the kinetochore that was imaged is highlighted with a red box. A slice of the resultant tomogram is overlaid with the fluorescent signal at that position, demonstrating the accuracy of CLEM. In all cases, we identify the kinetochore at the area of CENP-C fluorescence. The location of the kinetochore locus within the tomogram is shown by the blue dashed box and expanded in the inset. Each kinetochore inset is colored according to its morphology as defined in Figure 5E. In this dataset, tomograms colored red (Total loss of distinct architecture: Tomograms 3 and 6) also show reduced-size kinetochore complexes but are scored in this way since the kinetochore components are disorganized within the chromatin. Densities corresponding to disorganized components of the fibrous corona are present in all tomograms shown. Scale bars, 50 nm. **C.** Kinetochore architecture is preserved in CENP-N-eGFP-AID cells without CENP-N degradation as shown in example tomograms from the CENP-N-eGFP-AID -IAA condition. In this condition CENP-N is not degraded, and as expected the size of kinetochore complexes and kinetochore architecture match that seen in our initial preparation from CENP-A-rsEGFP2 cells (Fig. 1, 2, S2, S3). Tomogram slices are shown as in (B) except that fluorescent foci are yellow due to colocalization of CENP-C-cy3 and CENP-N-eGFP.

**Figure S10:**
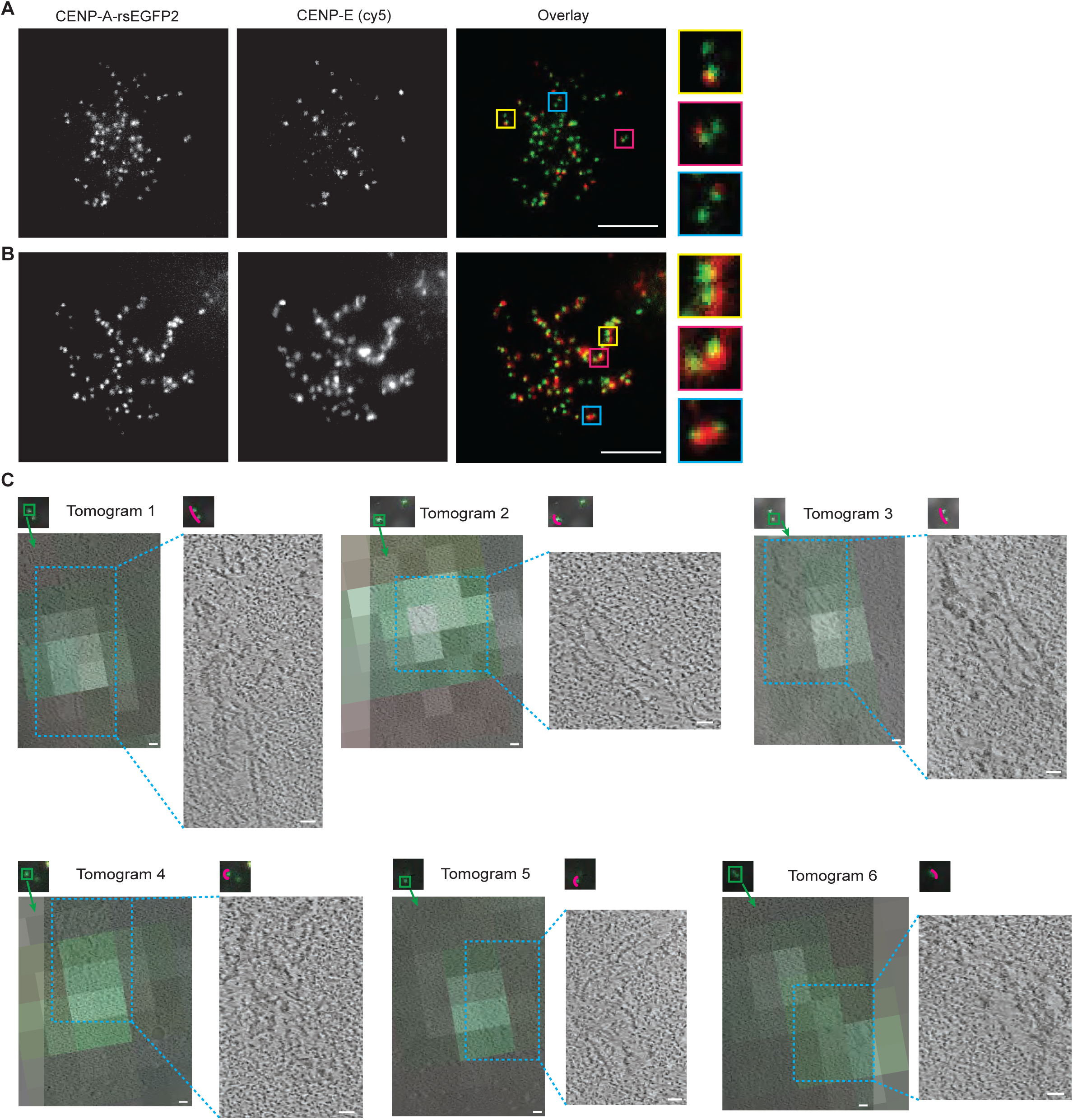
Prolonged detachment from the mitotic spindle leads to expansion of the fibrous corona. A. Immunofluorescence images from CENP-A-rsEGFP2 cells treated with 16 hours of STLC and 30 min nocodazole (original preparation: as in Figure 1a) with immunolabeling of the corona component CENP-E. A representative cell is shown, accompanied by expanded images of paired kinetochores. Chromosomes under this condition experience only brief detachment from the mitotic spindle and so kinetochores assemble either no corona or only a small corona as demonstrated by CENP-E fluorescence. Scale bars, 5 μm. **B.** Immunofluorescence images from CENP-A-rsEGFP2 cells treated with 16 hours nocodazole (as in Fig. 7A, lower panel) with immunolabeling of CENP-E. A representative cell is shown, accompanied by expanded images of paired kinetochores. Under this condition chromosomes experience prolonged detachment from the mitotic spindle due to microtubule depolymerization by nocodazole and so kinetochores assemble large, extended coronas as seen by the distribution of CENP-E fluorescence in these images. **C.** Further examples of expanded fibrous coronas seen in tomograms from the preparation using CENP-A-rsEGFP2 cells with prolonged nocodazole treatment (Fig. 7A, lower panel). The tomograms in this condition show longer fibrils forming an extended meshwork that can extend more than 1 μm in length. In each case the “double dot” CENP-A fluorescence pattern of a mitotic chromosome is shown and the kinetochore that was imaged is highlighted with a green box. A slice of the resultant tomogram is overlaid with the fluorescent signal at that position, demonstrating the accuracy of CLEM. The location of the fibrous corona within the tomogram is shown by the blue dashed box and expanded in the inset. The trajectory of the corona fibrils shown relative to the sister kinetochores of the chromosome imaged are shown in pink. In Tomograms 1, 3, and 6, the corona fibrils are seen to bridge across both sister kinetochores, whereas in Tomograms 2, 4, and 5 the corona fibrils wrap around a single kinetochore, both of which are consistent with fibrous corona profiles seen in mitotic cells as shown in (B). Tomogram 6 captures a fibrous corona spanning both sister kinetochores of a chromosome. As in Figure S3C, this chromosome was oriented on the grid such that the sister kinetochores were near each other in the x-y plane. Scale bars, 50 nm.

## Supplementary Information

Video S1: Tomogram of kinetochore from CENP-A-rsEGFP2 cell line shows kinetochore complexes within a single chromatin clearing.

This video begins above the surface of the chromosome, and kinetochore complexes are visible at the very surface of the chromosome. The video pans through the full kinetochore area and into a region of ordinary mitotic chromatin, then reverses back through the kinetochore area, building up a model of this locus. Kinetochore complexes are shown in cyan, and the chromatin clearing is outlined in red. The video also notes non-biological features of the tomogram including elements of the EM grid carbon film and erased gold fiducials. Scale bar, 20 nm.

Video S2: Tomogram of kinetochore from CENP-A-rsEGFP2 cell line shows a kinetochore composed of two chromatin clearings coalescing at the chromosome surface.

This video begins in ordinary mitotic chromatin and then pans into a kinetochore area in which kinetochore complexes can be seen within two distinct chromatin clearings. At the surface of the chromosome, these two clearings coalesce into one. The video then reverses back through the kinetochore area, building up a model of this locus. Kinetochore complexes are shown in cyan, and the chromatin clearing is outlined in red. The video also notes non-biological features of the tomogram including elements of the EM grid carbon film and erased gold fiducials. Scale bar, 50 nm.

Video S3: Details of individual kinetochore complexes, multimers thereof, and connections between adjacent complexes.

This video shows the same tomogram as in Video S1, but with additional detail of kinetochore complexes as shown in Figures 3 and S6. The video highlights an example individual kinetochore complex shown from multiple angles and density corresponding to a nucleosome within this complex. It also pans through multimers of kinetochore complexes and their associated nucleosomes. Finally, multiple instances of connections between adjacent kinetochore complexes are highlighted. The video also notes non-biological features of the tomogram including elements of the EM grid carbon film and erased gold fiducials.

Video S4: Tomogram of kinetochore from CENP-C-AID-eYFP cell line after CENP-C removal shows loss of kinetochore architecture.

This video shows a tomogram from the CENP-C-AID-eYFP cell line in the +IAA condition where CENP-C is degraded prior to chromosome isolation. The architecture of the kinetochore is greatly perturbed in this tomogram. The kinetochore area can be distinguished from surrounding chromatin, but the kinetochore complexes are indistinct and the boundary between the kinetochore and surrounding chromatin is poorly defined. This video pans from one edge of the chromosome to the other, and then reverses. The video also notes non-biological features of the tomogram including elements of the EM grid carbon film and erased gold fiducials. Scale bar, 50 nm.

Video S5: Tomogram of kinetochore from CENP-N-eGFP-AID cell line after CENP-N removal shows disruption of inner kinetochore complexes but retained chromatin clearing.

This video shows a tomogram from the CENP-N-eGFP-AID cell line in the +IAA condition where CENP-N is degraded prior to chromosome isolation. The inner kinetochore complexes are notably reduced in size but surrounded by an intact chromatin clearing. Disorganized fibrous corona components are also visible at the kinetochore. Scale bar, 50 nm.

Video S6: Tomogram of kinetochore from CENP-A-rsEGFP2 cell line after prolonged spindle perturbation shows expansion of the fibrous corona.

This video shows a tomogram in which the fibrous corona is greatly expanded due to prolonged detachment from the mitotic spindle prior to chromosome isolation. The video starts at the chromosome edge (at the EM grid carbon film), pans through the full corona/kinetochore area into an area of ordinary mitotic chromatin, and then reverses through the kinetochore area and builds up a model of the corona fibrils. The video also notes non-biological features of the tomogram including elements of the EM grid carbon film and erased gold fiducials. Scale bar, 50 nm.

Video S7: Tomogram of kinetochore from CENP-C-AID-eYFP cell line after CENP-C removal, with kinetochore over a hole in the EM grid carbon film.

This video shows a tomogram from the CENP-C-AID-eYFP cell line in the +IAA condition where CENP-C is degraded prior to chromosome isolation. As in Video S4, the architecture of the kinetochore is greatly perturbed in this tomogram, but in this case the kinetochore is located over a hole in the EM grid carbon film. Scale bar, 50 nm.

## Notes

### Competing Interest Statement

The authors have declared no competing interest.

